# Efficacy and safety of glycosphingolipid SSEA-4 targeting CAR-T cells in an ovarian carcinoma model

**DOI:** 10.1101/2022.09.02.506335

**Authors:** HJ Monzo, M Hyytiäinen, E Elbasani, K Kalander, J Wall, L Moyano-Galceran, J Tanjore-Ramanathan, J Jukonen, P Laakkonen, A Ristimäki, JW Carlson, K Lehti, S Salehi, P Puolakkainen, C Haglund, H Seppänen, S Leppä, PM Ojala

## Abstract

Chimeric antigen receptor (CAR) T-cell immunotherapies for solid tumors face critical challenges such as heterogeneous antigen expression. We characterized SSEA-4 cell-surface glycolipid as a target for CAR-T cell therapy. SSEA-4 is mainly expressed during embryogenesis but is also found in several cancer types making it an attractive tumor-associated antigen. Anti-SSEA-4 CAR-T cells were generated and assessed pre-clinically *in vitro* and *in vivo* for anti-tumor response and safety. SSEA-4 CAR-T cells effectively eliminated SSEA-4 positive cells in all tested cancer cell lines whereas SSEA-4 negative cells lines were not targeted. *In vivo* efficacy and safety studies using NSG mice and the high-grade serous ovarian cancer cell line OVCAR4 demonstrated a remarkable and specific anti-tumor response at all CAR-T cell doses used. At high T cell doses, CAR-T cell-treated mice showed signs of health deterioration after a follow-up period. However, severity of toxicity was reduced with delayed onset when lower CAR-T cell doses were used. Our data demonstrate the efficacy of anti-SSEA-4 CAR-T therapy; however, safety strategies, such as dose-limiting and/or equipping CAR-T cells with combinatorial antigen recognition should be implemented for its potential clinical translation.

## 1. Introduction

Cancer immunotherapies aimed at harnessing the immune system to fight refractory malignancies are rapidly evolving and have a promising potential. Based on the cancer immunosurveillance hypothesis [1], chimeric antigen receptor T (CAR-T) cell therapies targeting CD19 [2] have shown remarkable success in acute lymphoblastic and B-cell lymphomas [3]. In CAR-T cell therapy, patients’ T cells are genetically engineered *ex vivo* to express synthetic single-chain variable fragment (scFv) chimeric antigen receptors (CAR) against tumor antigens and subsequently infused back into patients, mounting a cytotoxic anti-tumor response upon antigen recognition [4].

In contrast to hematological malignancies, CAR-T cell therapies for solid tumors have produced limited efficacy and currently face multifactorial challenges for successful clinical translation. Solid tumors grow in functional association with the tumor microenvironment (TME) [5]. Typically composed of immune and stromal cells, blood vessels, and extracellular matrix (ECM), the TME constitutes a physical barrier and hostile immunosuppressive environment that CAR-T cells must overcome to target cancer cells [6, 7]. Furthermore, solid tumors are heterogeneous and express a diversity of tumor-associated antigens (TAA), which impair single-antigen CAR-T cell efficacy and lead to antigen escape and tumor relapse. Addressing these challenges in CAR-T cell therapy for solid tumors is thus necessary, and innovative strategies, such as the identification of actionable tumor antigens and cancer stem-like cell (CSC) targeting, are currently under active investigation [8-10].

CAR-T cells can trigger life-threatening toxicities, including cytokine release syndrome, neurotoxicity, and on-target off-tumor activity due to TAA expression in normal tissues [11]. Management strategies for CAR-T cell therapy toxicities involve systemic immunosuppression, CAR-T cell depletion using antibody-dependent cytotoxicity, and CAR-T cell dose evaluation to prevent potential off-targeting [12-14]. Analogous to normal stem cells (SCs), CSCs are a rare and persistent subpopulation of cancer cells with a strong ability to self-renew and produce heterogeneous progenitor cells that comprise the bulk of the tumor. CSCs have been identified in various malignancies, including ovarian, pancreatic, and gastric cancers [15-17], and are considered drivers of tumorigenesis, metastasis, and cancer progression owing to their therapeutic resistance. Accordingly, CSCs constitute a potential key target for CAR-T cell therapy [18, 19].

Stage-specific embryonic antigen-4 (SSEA-4) is a developmentally regulated glycosphingolipid (GSL). SSEA-4 expression has been reported to be restricted mainly to early embryogenic stages and pluripotent SCs and is mostly absent in differentiated cells and tissues [20]. GSLs play essential roles in cell adhesion, proliferation, and signal transduction. Consistently, malignant transformation encompasses cellular GSL re-programming [21]. In functional association with tumorigenesis and cancer progression [22], SSEA-4 is expressed in ovarian, breast, lung, pancreatic, and brain cancer cells, as well as in breast and ovarian CSCs [23, 24], thereby making SSEA-4 an attractive target antigen for CAR-T cell therapy [25-27]. In this study, we analyzed the expression of SSEA-4 in different human tumor tissues, as well as in cultured ovarian, pancreatic, and gastric cancer cell lines, and tested SSEA-4-targeting CAR-T cells pre-clinically *in vitro* and *in vivo*, with ovarian cancer cells, for their therapeutic efficacy and safety.

### 2. Materials and methods

### 2.1. Tissue processing and staining

Fresh frozen HGSC tumor samples were embedded in Tissue-Tek OCT compound (Sakura Finetek), frozen, and cut into 8-μm cryosections. Frozen sections were placed in 4% PFA in PBS for 10 min and washed with PBS for 10 min. For immunofluorescence staining sections were blocked with TNB blocking buffer (0.1 M Tris–HCl, pH 7.5; 0.15 M NaCl; 0.5% (w/v) blocking reagent (PerkinElmer, Cat # FP1020) for 30 min at RT. The sections were incubated overnight at 4 °C in a humidified chamber with SSEA4 antibody (1:100, GTX48037, Genetex; SSEA-4 hexaose [Elicity #GLY131-90%] blocking glycan was used as a negative control at 100 µM). After washes with TNT (0.1 M Tris-Cl, pH 7.5; 0.15 M NaCl; 0.1% (v/v) Tween 20), sections were incubated with PAX8 antibody (1:250, #10336-1-AP, Proteintech) and anti-mouse Alexa Fluor Plus 555 secondary antibody in TNB for 90 min and washed with TNT. The sections were then incubated with anti-rabbit Alexa Fluor Plus 488 secondary antibody in TNB for 90 minutes, washed with TNT, and rinsed with PBS. Slides were mounted with VECTASHIELD Antifade Mounting Medium containing DAPI. Confocal micrographs of immunofluorescence staining were obtained using a confocal microscope (LSM 800) with a C-Apochromat 40×, 1.2 NA water objective lens, and a Plan-Apochromat 20×, 0.8 NA objective lens (Carl Zeiss). For immunohistochemistry, sections were blocked with 3% BSA in PBS for 30 min and incubated with SSEA-4 antibody (GTX48037, Genetex) overnight, followed by incubation with Brightvision poly HRP-anti-mouse antibody, washing, DAB solution, and hematoxylin staining using standard protocols.

### 2.2. Cancer cell culture

OVCAR4 and Caov3 (human ovarian adenocarcinoma; RRID:CVCL_1627 and LGC Nordic ATCC HTB-75, respectively), HPAC and MIA PaCa-2 (human pancreatic adenocarcinoma; LGC Nordic, ATCC CRL-2119, and CRL-1420, respectively), and MKN45 and MKN28 (human gastric adenocarcinoma; RRID:CVCL_0434 and RRID:CVCL_1416, respectively) were cultured in RPMI medium (Roswell Park Memorial Institute 1640. #BE12-167F, Gibco) supplemented with 10% fetal bovine serum (FBS, qualified Brazil. #10270106 Gibco), 2 mM L-glutamine (#BE17-605E, Gibco), 100 U/mL penicillin, and 100 μg/mL streptomycin (#DE17-602E, Gibco) and subcultured in phosphate-buffered saline (1X; PBS. #MT21040CV, Gibco) and trypsin-EDTA solution (#BE17-161E; Lonza, Basel, Switzerland). HEK293T cells (human embryonic kidney; LGC Nordic ATCC CRL-3216) were cultured in DMEM high glucose (#BE12-614Q, Lonza), supplemented with 2 mM L-glutamine, 10% FBS, 100 U/mL penicillin, and 100 μg/mL streptomycin, and sub-cultured with PBS and trypsin. All the cell lines were authenticated using the Genome Analysis Infrastructure (HiLIFE; Helsinki Institute of Life Science) and assayed at low passage numbers. All cells were incubated at 37°C in 5% CO2 and routinely tested for mycoplasma-free status (Eurofins Genomic Europe, Germany).

### 2.3. Flow cytometry analysis

0.5-1×10^6^ Cells were labeled per tube in staining buffer (1% BSA/2 mM EDTA/PBS). Cells were fixed with 4% PFA for 10 min (where appropriate) at room temperature, 3x washed in buffer (2 mM EDTA/PBS), blocked for 30 min in staining buffer, and subsequently stained with primary and secondary antibodies in 100 µL volume, with 3x washes in between where appropriate, and after staining. The cells were resuspended in 200 µL wash buffer and analyzed using BD Accuri C6 Flow Cytometer Analyzer and Accuri Cflow Plus software. The gating strategy followed FSC-A *vs* SSC-A size and granularity, FSC-H *vs* FSC-A doublets, and corresponding FL *vs* FSC-A marker detection. Negative controls included isotypes, not stained samples, and where appropriate NT-T cells were used. The following antibodies were used in this study: Alexa Fluor 488 anti-human SSEA-4 (#330412, clone MC-813-70, Mouse IgG3 κ, dilution 1:100, Biolegend); Alexa Fluor 647 anti-human SSEA-4 antibody (#330408, clone MC-813-70, Mouse IgG3 κ, dilution 1:100, Biolegend). Cetuximab (#A2000, Human IgG1, dilution 1:100, Selleckchem); Alexa fluor 488 Goat anti-Human IgG Secondary (#A-11013, dilution 1:100, Invitrogen); APC anti-human CD4 Antibody (#357408, Rat IgG2b κ, dilution 1:20, Biolegend); FITC anti-human CD8 Antibody (#344704, Mouse IgG1 κ, dilution 1:20, Biolegend); Alexa Fluor 647 anti-human CD3 Antibody (#300322, Mouse IgG2a κ, dilution 1:100, Biolegend); APC Rat IgG2a κ Isotype Ctrl (#400512, dilution 1:20, Biolegend); FITC Mouse IgG κ Isotype Ctrl (#00108, dilution 1:20, Biolegend); APC Mouse IgG2a κ Isotype Ctrl (#981906, dilution 1:100, Biolegend).

### 2.4. Immunofluorescence

OVCAR4 and Caov-3 cells were grown on glass coverslips at low confluence for 24 h. Next, the cells were fixed with 4% PFA (#HL96753.1000, without methanol, Histolab), blocked with staining buffer for 30 min, stained with Alexa Fluor 647 anti-human SSEA-4 antibody (dilution 1:100) overnight at 4ºC, 3x washed with PBS, and counterstained with Hoechst 33342 (#H21492, Invitrogen). The slides were mounted with mowiol (#81381; Sigma-Aldrich). Slides were visualized with Axio Imager (Zeiss upright epifluorescence microscope) with an EC Plan Neofluar 40x 1.3 NA water objective lens and Plan-Apochromat 63x 1.4 NA oil objective lens (Zeiss). Images were acquired using Hamamatsu Orca Flash 4.0 LT B&W camera for fluorescence and Zeiss Zen 2 software.

### 2.5. Luciferase/GFP expression

Mammalian gene expression lentiviral vector pLV-EF1A>Luc2:T2A:Puro:F2A:TurboGFP (#VB210127-1246ann, VectorBuilder) was used to produce luc2/tGFP carrying lentivirus. The expression vector and packaging plasmids, pLP1, pLP2, and pLP/VSVG, were transfected into HEK293T cells. The viral supernatant was collected after 72 h, 0.45µm filtered, and precipitated with PEG-it (#LV825A-1, System Biosciences). The virus precipitates were resuspended in PBS and stored at -80°C. Tumor cells were transduced with a MOI of 10 for 24 h using Vectofusin-1 (10 µg/mL; #130-111-163, Miltenyi Biotec), with the following selection in Puromycin 1-2 µg/ml (#P8833, Sigma-Aldrich).

### 2.6. Generation of anti-SSEA-4 and anti-GFP CAR vector

The anti-SSEA-4 scFv was designed by CHO Pharma USA Inc. and inserted into a 2^nd^ generation CAR lentivector containing the CD28z co-stimulatory/activation domains; EGFRt sequence was cloned downstream of the CD28z sequence (Creative Biolabs; US). For anti-GFP CAR-T cells, anti-SSEA-4 scFv was replaced with anti-GFP scFv (Creative Biolabs). The CMV promoter was replaced with EF-1α for optimal expression of long RNA encoding multiple gene products, and the AmpR cassette was replaced with KanR to fulfil regulatory requirements in pre-clinical assessment and clinical trials (FIMEA; Finnish Medicines Agency) (Oxford Genetics; UK). VIVE Biotech (ES) identified and removed a poly A-like sequence in EGFRt.

### 2.7. Generation of human anti-SSEA-4 CAR-T cells

Human blood samples (∼ 40 ml) were obtained from healthy donors at the Finnish Red Cross Blood Services (FRCBS) and subsequently diluted 1:4 in PBS. Peripheral blood mononuclear cells (PBMCs) were separated by density gradient centrifugation using Ficoll-Paque Premium (#17-5442-02; GE Healthcare). PBMCs were washed 3x with PBS, filtered with a 30 µm cell strainer (#43-50030-03, Pluriselect), and cryopreserved in cryotubes (#177280, Nunc) in LN2 at 20×10^6^ cells/ml in 10% DMSO (cell culture reagent, #02196055-CF, MPBio)-Albunorm (200 g/l, #054343, Octopharma). CAR-T cells were generated following the Miltenyi protocol using TexMACS medium (#130-097-196), human CD4+ and CD8+ T-cell isolation kits (#130-096-533 and #130-096-495, respectively), T Cell TransAct human (#130-111-160), human IL-7/IL-15 cytokines (#130-095-362 and #130-095-764, respectively), and Vectofusin-1 (#130-111-163, Miltenyi Biotec). Briefly, PBMCs were thawed in a 37°C water bath for a few minutes, washed 2x with TexMACS medium, and incubated in TexMACS for 3 h. Subsequently, CD4^+^ and CD8^+^ T cells were isolated, combined, and activated/expanded in TexMACS medium supplemented with TransAct, IL7 (500 IU/mL), and IL-15 (84 IU/mL). After 24 h, CD4+ and CD8+ T cells were transduced with anti-SSEA-4 CAR lentivirus with a MOI of 5 for 24 h, using Vectofusin-1 (10 µg/mL) and spinoculation (800 × g for 30 min at 32°C). Subsequently, the culture medium was replaced with TexMACS supplemented with cytokines, and the cells were cultured for an additional 8-10 days, with half of the media changed every 2-3 days. For anti-GFP (mock) CAR-T cells, CD4+ and CD8+ T cells were transduced with anti-GFP CAR lentivirus.

### 2.8. Cytotoxicity assays

Cytotoxic killing of target cells was assessed using Thermo Scientific Black 96-Well Immuno Plates. Tumor cells were plated at 0.5×104 (OVCAR4) and 1×10^4^ (Caov-3, HPAC, MIA Paca-2, MKN45, and MKN28) cells/well. All target cells expressed the Firefly luciferase (FLuc) and turboGFP (tGFP) reporter genes to facilitate detection. After overnight culture, anti-SSEA4, anti-GFP CAR-T, and NT-T cells were added at the indicated E:T (effector : target cell ratio), and incubated for 72 h. Cell viability was measured using the Steady-Glo Luciferase Assay System (Promega) as indicated by the manufacturer in FLUOstar Omega Filter-based multi-mode microplate reader (BMG LabTech). Tumor cell-killing efficacy is represented as % cell viability relative to untreated cultures (E:T=0).

### 2.9. CFSE-dilution assay T cell proliferation

CAR and NT-T cells were labeled with CFSE (#C34554, CellTrace CFSE Cell Proliferation Kit Molecular Probes) according to the manufacturer’s instructions and at 1µM working concentration. 2.5×10^5^ CAR or NT-T cells were incubated with SSEA-4 and SSEA-3 (#GLY131 and #GLY122, respectively, Elicityl-Oligotech) at 200 µ M for 48 h in 24 well plate (Nunc). 5×10^5^ CAR and NT-T cells CFSE-labeled were incubated with or without 1×10^6^ OVCAR4 and Caov-3 cells for 48 h in RPMI-10% FBS-Pen/Strep and L-Glut medium in 10 cm cell culture dishes (Nunc). CFSE dilution was analyzed using flow cytometry on CD3^+^-gated cells; isotype/unstained cultures were used as CD3-control gating, and CFSE-unlabeled CAR and NT-T cells were used as gates for negative CFSE detection. Cell proliferation was assessed in relation to the corresponding CFSE-labeled CAR and NT-T cells cultured in the absence of cancer cells.

### 2.10. Cytokine secretion in *in vitro* assays

Secretion of human IFNγ was assessed from the supernatants of cytotoxic assays where cancer and T cells were co-cultured for 72 h in a 24-well format (0.5×10^5^ CAR and NT-T cells co-cultured with 1×10^5^ OVCAR4 cells), and mouse plasma samples, using ELISA MAX Deluxe Set human (IFNγ #430104, IL-2 # 431804, and IL-6 #430504, BioLegend) according to the manufacturer’s instructions. The absorbance was measured at 450 nm using FLUOstar reader.

### 2.11. Animals and ethics statement

All mice were housed in the Laboratory Animal Center facility, Biomedicum 1, Helsinki University, under pathogen-free conditions and in ambient rooms with a 12 h light/dark cycle, filtered air at 22 ± 2°C temperature, and 55 ± 15% relative humidity. Mice were caged in groups of two or three per IVC cage (Green-Line mice. Emerald. Tecniplast) on autoclaved ASPEN wood chip bedding, with ad libitum access to laboratory chow (Tekland Global Rodent Diet; T2916CMI.16) and sterile water. Animal experiments in this study were approved by the Finnish National Animal Experiment Board (Eläinkoelautakunta ELLA. Approved license: ESAVI/10548/2019) and conducted in accordance with the Finnish Act on the Protection of Animals Used for Scientific or Educational Purposes, and the University of Helsinki guidelines.

### 2.12. Animal studies and bioluminescence imaging

Female NSG NOD.Cg-Prkdc^scid^ Il2rg^tm1Wjl^/SzJ mice (4–5 weeks old) were purchased from The Jackson Laboratory (Scanbur). 2×10^6^ OVCAR4 Fluc-tGFP cells, in 100 µL PBS, were injected intraperitoneally into isoflurane-anesthetized mice (Attane vet, 100 mg/g) and grown for 13 days. Mice were randomized for similar tumor load per mice in all groups. Subsequently, CAR and NT-T cells were infused intravenously (d0) at the specified T-cell dose in 100 µ l PBS. *In vivo* CAR-T cell dose is defined in this study as the specific number of CAR-T cells and was calculated based on the transduction percentage. Equal numbers of NT-T cells were used for each cell dose (Supplementary Fig. S2 and S5). For the no-treatment control group, mice were infused with 100 µL of PBS. Tumor progression was monitored by BLI with CoA D-luciferin sodium salt substrate (#bc218, Synchem) injected intraperitoneally at 150 mg/kg. Bioluminescence images were acquired 10 min after D-luciferin injection with the SPECTRAL Lago X Imaging System and analyzed with Aura Imaging Software v3.1.0 (Spectral Instruments Imaging). Mice were monitored at regular intervals for behavior, activity, feeding, and weight. At the end of the experiment, the mice were anesthetized with ketamine-xylazine (#511485, Intervet; #148999, Orion Pharma) and blood samples were collected via cardiac puncture in EDTA capillary tubes (#076012, Kabe Labortechnik). Plasma and blood cells were separated by centrifugation (500 × g for 5 min at 4°C) and stored at -80°C. Organs/tissues were collected post-mortem, imaged for bioluminescence, and fixed/stored in 4% PFA (without methanol) at 4°C.

### 2.13. Cytokine secretion in *in vivo* assays

Human cytokine secretion was assessed in mouse plasma samples using the Human XL Cytokine Luminex Performance Panel Premixed Kit (#FCSTM18, R&D Systems), according to the manufacturer’s instructions.

### 2.14. Histopathology

Tissue samples were shipped in 10% formalin solution at RT to HistoWiz Inc. (US) for processing and histological examination by a certified pathologist.

### 2.15. Statistics

*In vitro* experiments were independently performed at least three times, and at least three mice per donor were compiled for *in vivo* experiments unless otherwise specified in the figure legends. Data processing was performed using Microsoft Excel 2016, and graphing and statistical analyses were performed using GraphPad Prism 9.0 Windows 64-bit software (San Diego, CA, USA). Data are reported as mean ± standard deviation, unless otherwise specified. Statistical analysis was performed using one-way analysis of variance with Tukey’s correction for multiple comparisons as indicated in the figure legends. Statistical significance was set at *p* < 0.05. *p*-values are indicated in the figure legends.

## 3. Results

### 3.1. Characterization of SSEA-4 expression in human cancer specimens and cell lines

SSEA-4 expression has been demonstrated in various types of cancer [28]. To corroborate and extend previous findings, we stained formalin-fixed paraffin-embedded (FFPE) clinical gastric cancer samples. We observed only a low signal or complete absence of SSEA-4 immunoreactivity in all studied samples (Supplementary Fig. S1A). Next, we investigated whether the lack of specific staining was due to the organic solvent treatment normally used during staining of FFPE samples. In fresh frozen (FF) tumor samples, we detected strong SSEA-4 immunoreactivity in several cancer types, including gastric cancer, (Fig. 1A-C). In contrast, the treatment of FF tumor tissue sections with a standard xylene/ethanol series resulted in SSEA-4 signal loss (Supplementary Fig. S1B). This finding indicates that the organic solvent treatment is not suitable for SSEA-4 detection. To confirm the specificity of the anti-SSEA-4 antibody, we stained FF pancreatic cancer samples with soluble SSEA-4 hexaose (blocking glycan). This resulted in the loss of the SSEA-4 signal, which was specific for SSEA-4, as the closely related glycolipid, SSEA-3, did not block SSEA-4 immunoreactivity (Supplementary Fig. S1C). SSEA-4 expression was prominent but heterogeneous in FF tissues from ovarian (OC), pancreatic (PC), and gastric (GC) cancers (Fig. 1A-C). In OC, 20 of 23 samples (87%) expressed SSEA-4 with variable intensity and extent (Fig. 1A). In some samples, SSEA-4 expression was observed in both tumor cells and the surrounding stroma, whereas in others, its expression appeared to be restricted to the stroma. Focusing on OC, and for a more precise analysis, we performed double immunofluorescence staining for SSEA-4 and tumor cell-specific PAX8, which revealed the localization of SSEA-4 in tumor cells or stroma (Fig. 1A). In PC, 19 of 23 samples (83%) expressed SSEA-4 (Fig. 1B). Tumor cell and acinar cell immunoreactivity were also detected. Similar to OC cases, some samples showed SSEA-4 signals only in the stroma. In GC, eight of the nine samples (89%) were positive for SSEA-4, with expression mostly localized to tumor cells (Fig. 1C).

**Figure 1.**
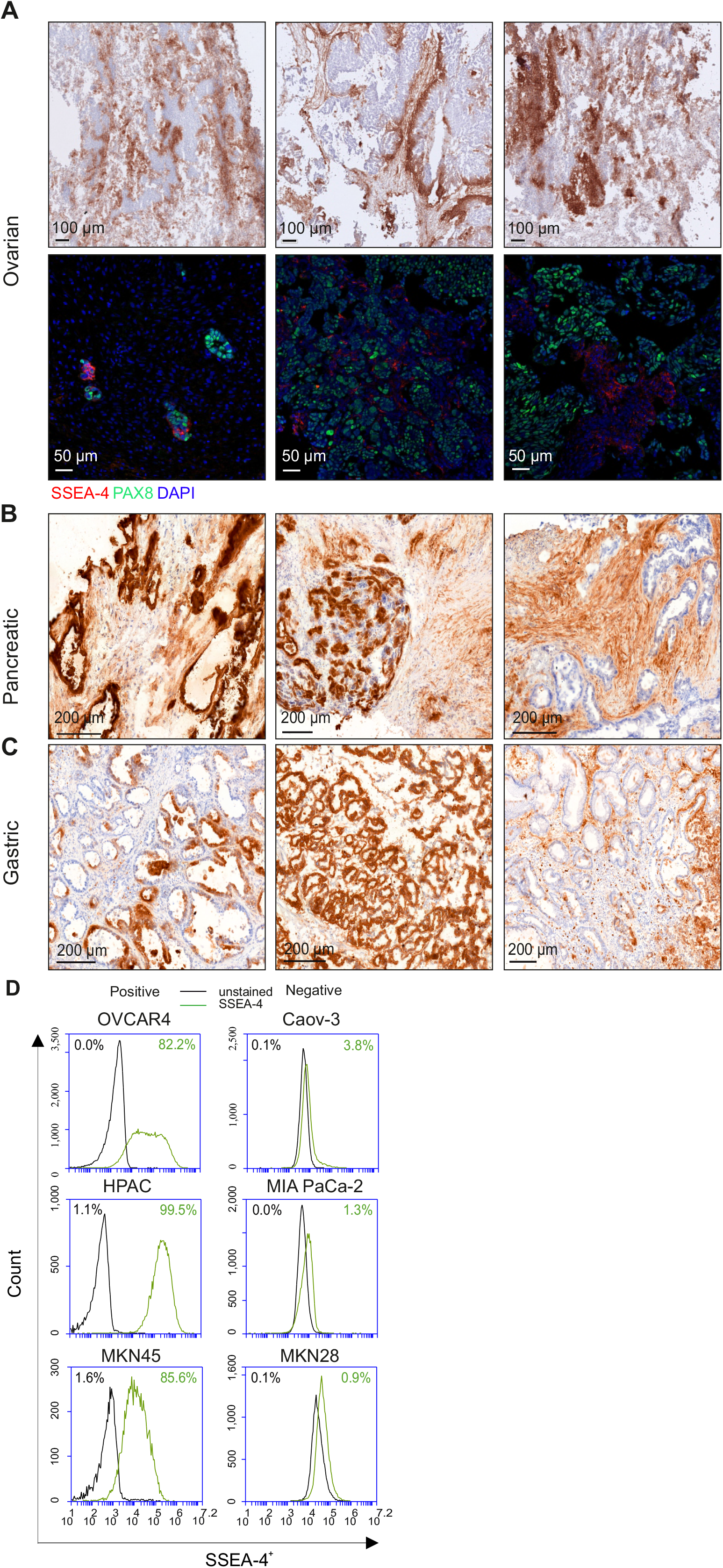
Analysis of SSEA-4 expression in tissues and cancer cell lines. **A-C**. SSEA-4 immunohistochemistry in (A) Ovarian, (B) pancreatic, and (C) gastric cancer patient samples. Ovarian cancer samples were also immunofluorescently stained with antibodies against SSEA-4 (red) and PAX8 (green) (A. lower panel). Representative samples are shown. **D**. Flow cytometry plots show SSEA-4 expression in human cancer cell lines. OVCAR4 (ovarian), HPAC (pancreatic) and MKN45 (gastric) were positive for SSEA-4, and Caov-3 (ovarian), MIA PaCa-2 (pancreatic) and MKN28 (gastric) were negative for SSEA-4. Unstained/isotype antibody-stained cells were used for gating the SSEA-4 negative cells.

To find suitable SSEA-4 expressing cell models, we analyzed SSEA-4 expression in a variety of cancer cell lines using FACS. Some cell lines representing the aforementioned cancer types were either highly positive (OVCAR4, HPAC, and MKN45; OC, PC, and GC, respectively) or very low/negative (Caov-3, MIA PaCa-2, and MKN28; OC, PC, and GC, respectively) for SSEA-4 (Fig. 1D). The FACS results were further confirmed by immunofluorescence staining of the OC cell lines OVCAR4 and Caov-3 (Supplementary Fig. S1D).

### 3.2. Generation and *in vitro* characterization of SSEA-4 specific CAR-T cells

To obtain optimal expression of a long RNA encoding multiple gene products and to fulfill regulatory requirements in the putative future clinical trials, a previously described anti-SSEA-4 CAR vector [29] was modified by replacing the cytomegalovirus (CMV) promoter with EF-1α (human elongation factor-1 alpha) and ampicillin (AmpR) cassette with kanamycin (KanR). For T cell transduction assessment, a truncated human epidermal growth factor receptor (EGFRt) [30] (Fig. 2A) was inserted downstream of the CD28 and CD3zeta sequences (costimulatory/activation domains). A poly A-like sequence in EGFRt was also removed. Anti-SSEA-4 CAR-T cells (hereinafter referred to as CAR-T) were generated from healthy donor-derived CD4^+^ and CD8^+^ T cell populations, transduced with a 60% average efficiency (Fig. 2B), and expanded for 10-12 days. CD4^+^/CD8^+^ quantification after expansion showing a slightly enhanced proliferation for CD8^+^ cells (Fig. 2C). Non-transduced T-cells from the same donor (NT-T) were used as control.

**Figure 2.**
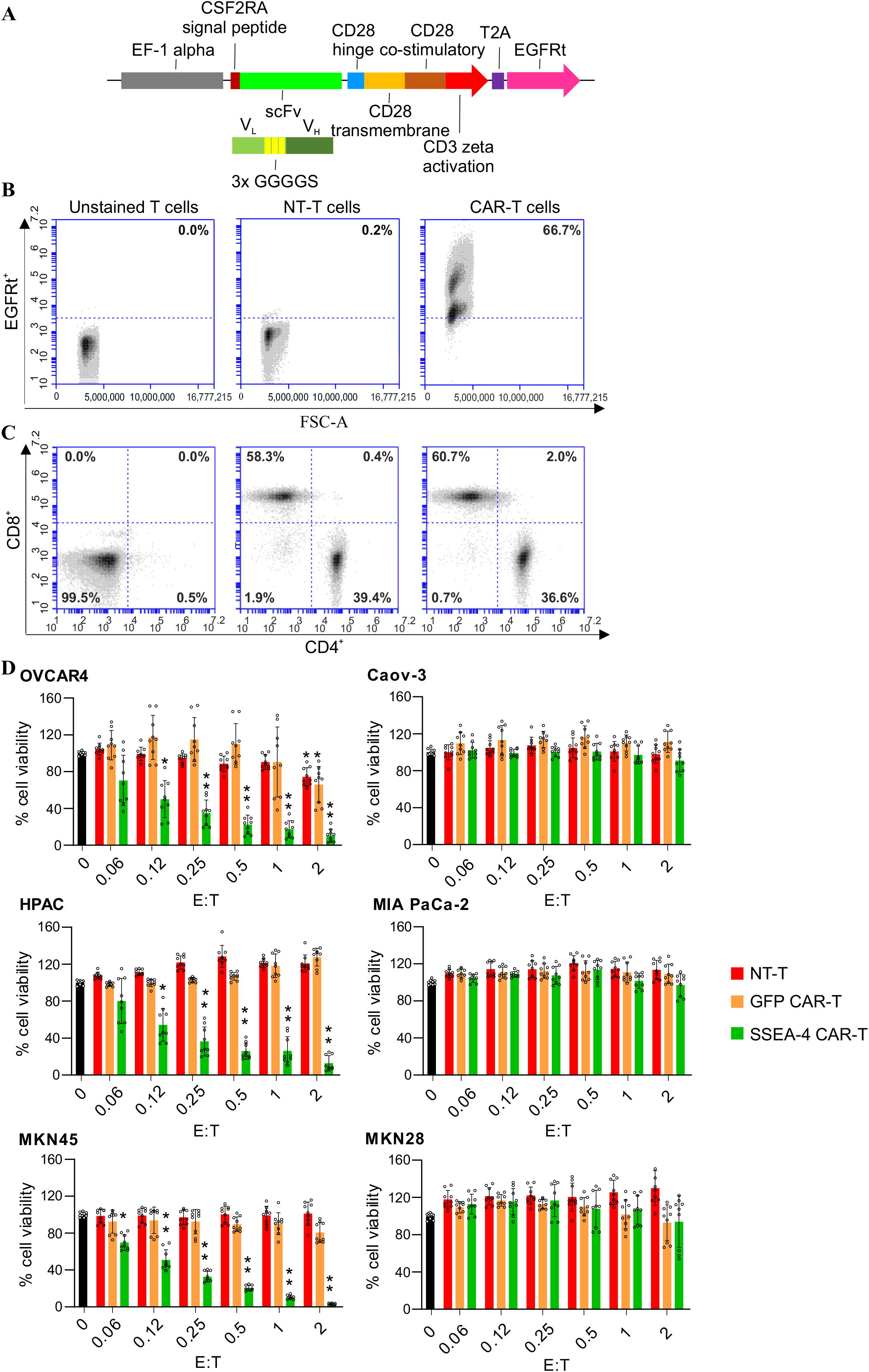
Anti-SSEA-4 CAR design, T cell transduction, and CAR-T cell anti-tumor response*in vitro*. **A**. Schematic representation of the SSEA-4 CAR lentivector. EF1-a promoter driving the expression of CSF2RA (colony stimulating factor receptor alpha 2) signal peptide followed by anti-SSEA-4 V_L_-(GGGGS)3-V_H_ scFv linked by the CD28 hinge to CD28-CD3z co-stimulatory and signaling domains. EGFRt gene is located downstream and separated by the T2A self-cleaving peptide. **B**. Flow cytometry plots showing percentage of transduced T cells assessed by detection of EGFRt expression using cetuximab antibody. Unstained T cells were used for gating the EGFRt negative cells and non-transduced (NT) T cells as a control. **C**. Flow cytometry plots showing the proportion of CD4^+^/CD8^+^ T cells in the cultures 10-12 days post transduction. Unstained/isotype T cells were used for gating the CD4/CD8 negative cells and NT-T as a control. **D**. *In vitro* cytotoxicity assays using co-cultures of SSEA-4 CAR-T, NT-T, and anti-GFP CAR-T cells at the indicated doses with SSEA-4^+^ (left panel; OVCAR4, HPAC, MKN45) and SSEA-4-(Caov-3, MIA PaCa-2 and MKN28; right panel) cancer cells for 72 h. Cancer cell killing efficacy was assessed by luminescence emission from the cancer cells, and is represented on the y-axis as the percentage of cell viability relative to untreated (E:T, effector : target cell ratio = 0) cultures. N=3. Mean ± SD. One-way ANOVA with Tukey’s multiple comparison tests. (**F**) * Indicates significant differences with T cell dose = 0. \*\**p=0*.*0001; *p*≤*0*.*03*. Not significant differences were attributed to *p*≥*0*.*05*.

To assess CAR-T anti-tumor response and specificity, we performed *in vitro* cytotoxicity assays in SSEA-4^+^ OVCAR4 (OC), HPAC (PC), and MKN45 (GC), as well as in SSEA-4-Caov-3 (OC), MIA PaCa-2 (PC), and MKN28 (GC) human cell lines. The specificity and efficacy of CAR-T cells were evaluated at different T-cell doses in comparison with NT-T and GFP (mock) CAR-T cells. CAR-T cells killed SSEA-4^+^ OC, PC, and GC cells efficiently (Fig. 2D) but did not show any significant cytotoxic activity against SSEA-4-cells. NT-T and GFP CAR-T cells showed no effect on any cell type tested, further corroborating the specific SSEA-4 targeted anti-tumor response of the CAR-T cells. The CAR-T cell response was dose-dependent and detectable at E:Ts (effector : target cell ratios) as low as 0.12 in all SSEA-4^+^ cell lines tested. In contrast to NT-T cells, the interaction of CAR-T cells with OVCAR4 cells, but not with Caov-3 cells, induced T-cell proliferation (Fig. 3A-C), indicating T-cell activation in response to SSEA-4. A significant increase in IFNγ and IL2 secretion was also observed in CAR-T cells, but not in NT-T cells, when co-cultured with OVCAR4 cells (Fig. 3D and E), further supporting the specific SSEA-4 -induced functional activation of CAR-T cells.

**Figure 3.**
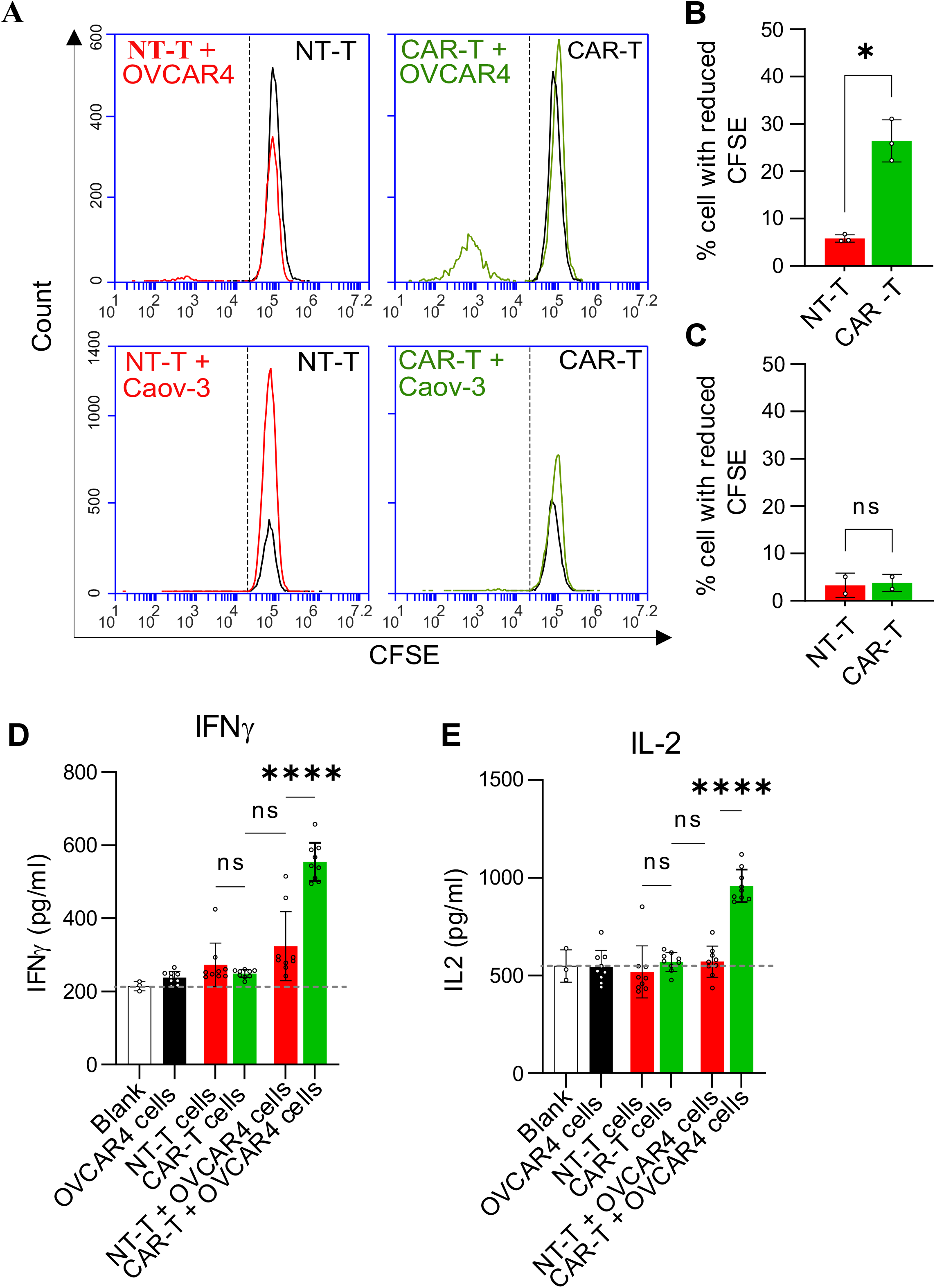
Characterization of anti-SSEA-4 CAR-T cells *in vitro*. **A**. Flow cytometry histograms of CFSE (Carboxyfluorescein succinimidyl ester) cell proliferation assays showing CAR-T cell proliferation upon interaction with SSEA-4^+^ OVCAR4 cells in contrast to SSEA-4-Caov-3 after 48 h. NT-T cells showed no proliferation in OVCAR4 and Caov-3 co-cultures. **B** and **C**. CFSE quantification of cells in A. **D** and **E**. Levels of IFNγ (**D**) and IL-2 (**E**) detected by ELISA from the supernatants of OVCAR4 +/-NT-T or SSEA-4 CAR-T cells at 72 h. (**B** and **C**) N=2. (**D-F**) N=3. Mean ± SD. One-way ANOVA with Tukey’s multiple comparison tests. (**B**) \**p=0*.*019;* (**D-F**) \*\*\*\**p=0*.*0001* and \*\*\**p=0*.*001*. Not significant differences, n.s., were attributed to *p*≥ *0*.*05*.

### 3.3. SSEA-4 CAR-T cells produce robust anti-tumor response in the OVCAR4 xenograft model

Next, we evaluated the *in vivo* efficacy of SSEA-4 CAR-T cells in ovarian cancer cell xenografts. NSG (NOD scid gamma) female mice were implanted intraperitoneally (i. p.) with 2×106 luciferase- and GFP-expressing OVCAR4 cells. After 13 days, mice were infused intravenously with T cell doses of 1×10^6^, 2×10^6^ and 3×10^6^ CAR-T cells (hereinafter referred as 1M, 2M and 3M), corresponding to 2.9×10^6^, 5.7×10^6^ and 8.6×10^6^ of total T cells based on 35% lentivirus transduction efficiency in the cell batch used in this experiment. Control mouse groups were infused with PBS or an NT-T cell dose equal to the total T cell number in the CAR-T groups (Supplementary Fig. S2A). Thirty-two days after T-cell infusion, CAR-T cell-treated mice in all dose groups presented a remarkable and specific reduction in tumor burden, in contrast to PBS- and NT-T cell-treated mice showing continuous tumor growth (Fig. 4A-D). However, mice treated with the 3M dose of NT-T cells displayed a similar reduction in tumor burden as the CAR-T cell-treated group, revealing an upper threshold of ≈ 5-6×10^6^ total T cells for the CAR-T specific anti-tumor response in the OVCAR4 i. p. xenograft model (Fig. 4D). All mice remained in apparently good body scoring conditions (BCS ≈ 3) during the 32-day experiment, except for 3M CAR-T mice, which showed a trend for weight loss at the later stages (Supplementary Fig. S2B). T cell activation was confirmed by measuring IFNγ levels in post-mortem mouse blood samples (Supplementary Fig. S3A). IFNγ secretion was significantly elevated in mice treated with 1M, 2M and 3M CAR-T cell doses compared to mice infused with PBS or NT-T cells. In line with the anti-tumor response of NT-T cells at high doses, elevated levels of IFNγ were also detected in 3M NT-T cell-treated mice. Bioluminescence imaging (BLI) of the post-mortem organs corroborated the specific reduction in tumor burden in mice treated with 1M, 2M and 3M CAR-T doses, showing a substantially reduced signal in all organs examined when compared to PBS- and NT-T-treated mice (Supplementary Fig. S3B and C). The cancer cell signal in our OVCAR4 xenograft model was preferentially associated with the spleen, pancreas, ovaries, intestine, and stomach. Similar to the whole-body BLI results, mice treated with the 3M NT-T cell dose showed reduced organ BLI signal.

**Figure 4.**
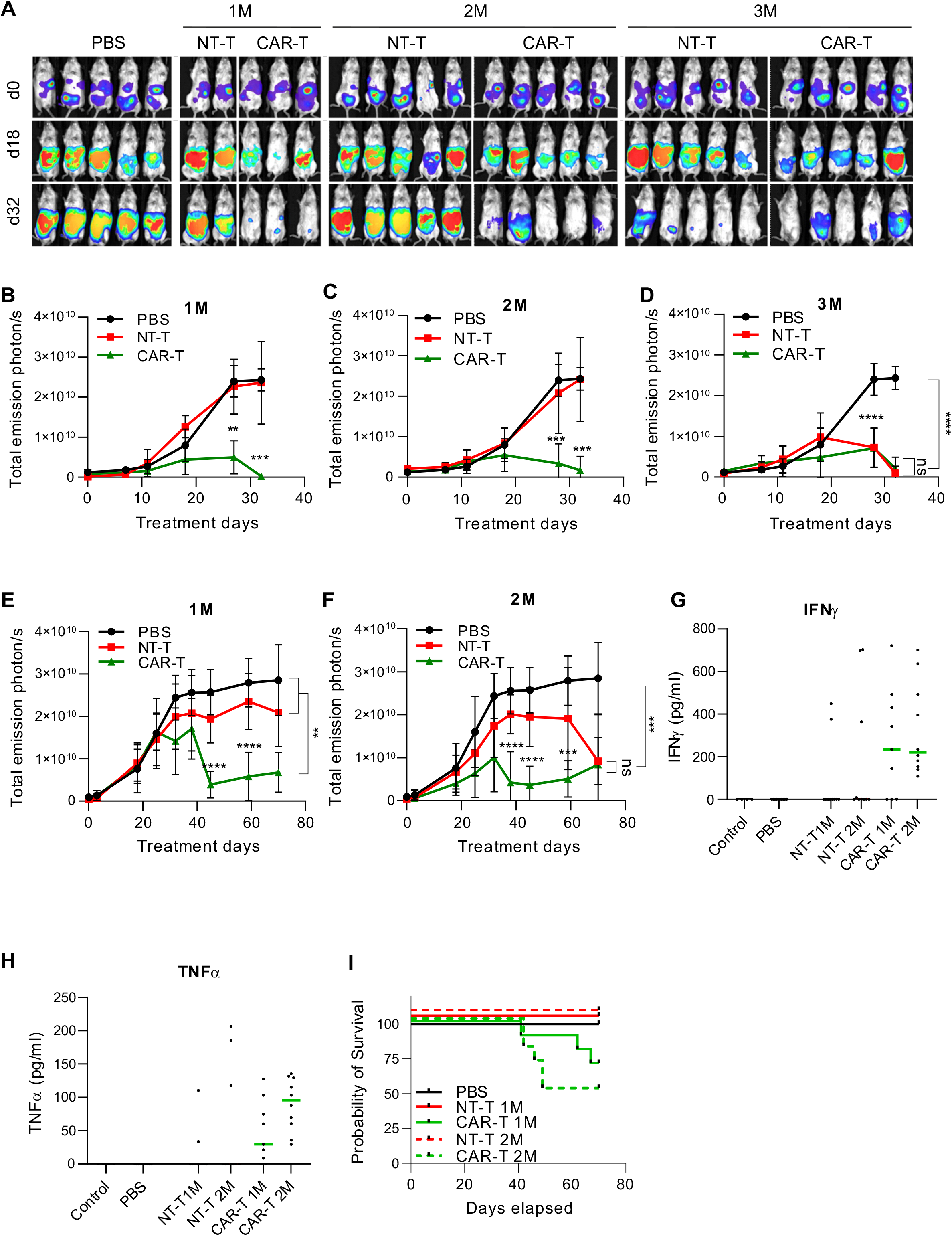
Anti-tumor response and survival in *in vivo* study. **A-D**. Short-term anti-tumor response of SSEA-4 CAR-T cells in NSG mice implanted intraperitoneally with 2×10^6^ FLuc-expressing OVCAR4 cells. (**A**) Representative BLI images of mice treated with 1M, 2M and 3M T cell doses at the indicated time points (days; d). **B-D**. BLI kinetics in the mice treated with SSEA-4 CAR-T at the indicated doses and compared to NT-T cells and PBS. N=5 for PBS, 2M and 3M groups. N=2 for NT-T 1M group. N=3 for 1M CAR-T group. Mean ± SD. One-way ANOVA with Tukey’s multiple comparison tests between PBS, NT-T and CAR-T groups at the indicated time points. (**B**) \*\**p=0*.*0018* and \*\*\**p=0*.*0004*, (**C**) *\*\*\*p=0*.*0009*, (**D**) *\*\*\*\*p=0*.*0001*. **E** and **F**. Long-term assay anti-tumor response of SSEA-4 CAR-T cells in NSG mice. (**E** and **F**) BLI kinetics in the mice treated with SSEA-4 CAR-T at the indicated doses and compared to NT-T cells and PBS. N=10 for all groups, except for NT-T 1M dose N=8. (**E**) *\*\*\*\*p=0*.*0001 and **p=0*.*0035*, (**F**) *\*\*\*\*p=0*.*0001* and *\*\*\*p=0*.*0004*. **G** and **H**. Quantification of IFNγ and TNFα using ELISA in the post-mortem mouse blood plasma samples of the long-term assay. N=10, except for control group (N=4), NT-T 1M (N=8), and CAR-T 1M group (N=9). **I**. Probability of survival for PBS and CAR/NT-T dose mice groups after 70-day treatment. (M=1×10^6^).

### 3.4. SSEA-4 CAR-T cells cause long-term adverse effects

Although there was no apparent toxicity in mice in the above-described 32-day efficacy *in vivo* study, modest signs of toxicity were observed in mice treated with the 3M CAR-T cell dose. We therefore carried out a long-term (70-day) safety, efficacy, and survival assay *in vivo* using 1M and 2M CAR-T cell doses, corresponding to 1.1×10^6^ and 2.2×10^6^ T cells based on 90% lentivirus transduction in the cell batch used in this experiment (Supplementary Fig. S4A). As previously described, tumor-bearing control mice were infused with PBS or an NT-T cell dose corresponding to the total number of T cells in the CAR-T groups. Tumor-free mice treated with a 1M dose were used to assess the potential toxicity of CAR/NT-T cells in the absence of OVCAR4 cells.

Corroborating the anti-tumor response of SSEA-4 CAR-T cells, both CAR-T doses led to a reduction in tumor burden compared to PBS treatment. While the 1M dose NT-T mice showed no significant reduction in tumor burden, the NT-T 2M dose produced a significant anti-tumor response at the later stages of the 70-day treatment (Fig. 4E and F, Supplementary Fig. S4B and C). Consistent with the anti-tumor activity of CAR-T cells, mice treated with both 1M and 2M doses showed elevated levels of IFNγ and TNFα (Fig. 4G and H). All mice remained in good health during the first 35-40 days of treatment. However, thereafter, some tumor-bearing CAR-T cell-treated mice showed weight loss, inactivity, and an overall low BCS (≈ 1-2). Three out of 10 mice with 1M dose, and 5 out of 10 mice with 2M dose, had to be sacrificed before the experimental endpoint (Supplementary Fig. S4B and C, and S5), translating in 70% and 50% survival rates, respectively (Fig. 4I). PBS, and the remaining CAR-T and NT-T mice were active and showed good overall condition (BCS ≈ 3), and normal activity without signs of health deterioration. Similarly, tumor-free mice treated with CAR-T and NT-T cells remained healthy and active during the 70-day study period (Supplementary Fig. S5). Altogether, our data demonstrated the specific anti-tumor response of SSEA-4 CAR-T cells, even at relatively low doses. At high doses, however, CAR-T cells produced apparent toxicities in mice, thus precluding their clinical translation at this stage.

### 3.5. SSEA-4 targeting of normal mouse cells as a potential cause for the *in vivo* adverse events during long-term SSEA-4 CAR-T cell treatment

To examine the possible on-target off-tumor effects in mice treated long-term with SSEA-4 CAR-T cells, we analyzed the expression of SSEA-4 in normal mouse tissues. Prominent SSEA-4 expression was detected in the kidneys, stomach, and intestine and, to a lesser extent, in the brain and heart. The SSEA-4 signal was sparse in the lungs, whereas the spleen, pancreas, and liver were mostly devoid of SSEA-4 (Supplementary Fig. S6).

Histopathological analysis of post-mortem tissues was performed on tissues with high (kidney and intestine) or moderate (lung) SSEA-4 expression. The analysis included the estimation of lymphocyte infiltration and tissue damage, each graded on a scale from 0 to 3. Analysis of the H&E-stained sections revealed perivascular/peribronchiolar lymphocyte infiltrates in the lungs, and in the intestinal wall of the CAR-T-treated mice compared to the NT-T-treated mice (average grading 0 *vs* 1.43, respectively for mice treated with 2M doses; Supplementary Fig. S7). Signs of aspiration pneumonia were observed in three CAR-T cell-treated and prematurely euthanized mice. However, there was no apparent, histologically appreciable, direct damage to the parenchymal cells (Supplementary Fig. S7).

To further corroborate the histopathological findings, we performed IF staining of total T cells (using anti-CD3) and specifically the CAR-T cells (anti-EGFRt) in the lungs, kidneys, and intestine (Supplementary Fig. S8). Consistent with H&E staining, the most prominent T cell deposits were observed in the lungs of both CAR- and NT-T cell-treated mice. CAR-T cells were also detected in the intestinal wall of CAR-T cell-treated mice. NT-T cells were not observed in NT-T-treated mice in the tissues analyzed.

## 4. Discussion

CAR-T cell therapy approaches in solid tumors face multifactorial challenges, including identification of actionable tumor antigens [11]. To tackle this obstacle, we conducted a pre-clinical CAR-T cell study using SSEA-4 as a TAA in ovarian, pancreatic and gastric cancer cell lines *in vitro* and assessed the efficacy and safety in an ovarian cancer xenograft model *in vivo*.

SSEA-4 has been shown to be expressed in pluripotent SCs, and developmentally regulated [20]. Consistent with its role in proliferation and migration, aberrant expression of SSEA-4 confers tumor cells with malignant stem-like traits such as proliferation, drug resistance, and metastasis [21, 31]. Thus, SSEA-4 reprogramming of tumor cells provides an attractive target for CAR-T therapies in solid tumors. In line with this, CAR-T cells targeting glycans such as MUC1 and GD2 have shown high therapeutic potential against triple-negative breast cancer and glioblastoma [32, 33]. Moreover, SSEA-4 CAR-T cells enable targeting of tumor cells with potential stem-like properties (CSCs), which are closely associated with tumor heterogeneity and relapse [19, 23, 34]. Clinical trials using CAR-T cells against CSC markers include CD133, CD171, and c-Met for ovarian, brain, and breast cancers and neuroblastoma [35].

Confirming previous results [28], SSEA-4 expression was detected in most of the cases of ovarian, pancreatic, and gastric cancer tissues. Based on the positive SSEA-4 expression in several cancer types, anti-SSEA-4 CAR-T cells were generated and functionally characterized for specific and effective anti-tumor responses.

SSEA-4 CAR-T cells showed specific and effective anti-tumor responses *in vitro* in SSEA-4^+^ ovarian, pancreatic, and gastric tumor cell lines. CAR-T cells produced elevated levels of IFNγ and IL-2 in killing assays with ovarian cancer cells and, corroborating the specific interaction with SSEA-4, CAR-T cell proliferation was induced in co-cultures with SSEA-4+ ovarian cancer cells. Current strategies to overcome CAR-T cell therapy challenges involve engineering CAR-T cells with powerful anti-tumor activity [11]. In this respect, SSEA-4 CAR-T cells demonstrated effective SSEA4+ targeting, even at very low E:T ratios in all *in vitro* cancer models.

High-grade serous ovarian carcinomas (HGSOC), represented by OVCAR4 cells in this study, constitute the most common and lethal gynecologic malignancies, typically forming immunologically “cold” tumors that are unresponsive to immune checkpoint blockage [36]. To date, responses to CAR-T therapies in HGSOCs have been relatively poor, mainly due to the deficient anti-tumor response of T cells and poor infiltration capacity in the tumor tissue [8]. In our *in vivo* OVCAR4 xenograft models, SSEA-4 CAR-T cells produced an effective anti-tumor response at dose levels ranging from 1M to 3M, thereby indicating that they could infiltrate into the tumor tissue and develop potent anti-tumor activity.

In a long-term safety assessment, SSEA-4 CAR-T cells produced severe adverse events in some mice, manifested by abnormal inactivity and progressive weight loss, indicating a dose-limiting toxicity for SSEA-4 CAR-T cells. Notably, in our OVCAR4 xenograft model, lower CAR-T cell doses produced an effective anti-tumor response, which was similar to that observed at higher CAR-T cell doses, but with a delayed onset and reduced severity of adverse effects. Moreover, mice treated with 1M dose of CAR-T cells showed moderate weight loss at the early stages of the treatment but progressively regained weight to normal levels and remained active and in good health condition (BCS ≈ 3) until the end of the study. The severity of the CAR-T cell associated adverse events is related, among the others, to CAR-T cell potency and dose [37]. Increasing the number of CAR-T cells has been shown to enhance tumor clearance but may also increase the risk of toxicities [38]. In line with this, strategies to prevent and manage toxicities include treatment with low or split doses of CAR-T cells [39, 40]. Based on our results, it is conceivable that, at very low doses or with a lower affinity scFv, SSEA-4 CAR-T cells might exert a potent anti-tumor response with significant lesser toxicity.

Targeting CSCs with CAR-T cell therapy constitutes a promising strategy for tumor eradication [35, 41]. However, the expression of CSC antigens may also occur in normal stem cells, thus increasing the risk of on-target off-tumor toxicities. In this regard, infiltrated SSEA-4 CAR-T cells in the SSEA-4 positive mouse intestinal wall may have impaired the normal absorption function of the gut, even though no direct damage to the epithelium was detected. Similarly, prominent lymphocyte infiltration was observed in the lungs of the CAR-T cell-treated mice. Accumulation of SSEA-4 CAR-T cells in the lungs and intestine might have produced severe inflammation, thereby affecting the normal respiratory and digestive functions in CAR-T-treated mice.

Tumor-free mice treated with CAR/NT-T cells at 1M dose showed no signs of health deterioration and remained active and healthy throughout the study. The absence of toxicities in tumor-free mice strongly suggests that in tumor-bearing mice an OVCAR4-dependent, over-stimulated SSEA-4 CAR-T cell immune response caused serious adverse effects. These findings are consistent with a recent study [29] reporting SSEA-4 CAR-T cell anti-tumor response in a pancreatic cancer mouse xenograft model, but with toxicities at the high CAR-T cell doses (Supplementary Figure 8 in [29]). NT-T cells at 3M dose, and to some extent at 2M dose, also reduced OVCAR4 tumor burden, thereby demonstrating a CAR-independent anti-OVCAR4 response. Further supporting this notion, our *in vitro* results showed a dose-dependent anti-OVCAR4 response in NT-T cell cultures. Taken together, these data indicate that OVCAR4 cells may be particularly prone to elicit CAR-independent anti-tumor T cell reactivity, i.e. graft-versus-tumor effect [42]. Accordingly, at high CAR-T cell doses, the occurrence of both CAR-dependent and CAR-independent interactions with OVCAR4 tumor tissues could have resulted in over-stimulated T-cell activation, inflammation and, consequently, damage to normal tissues, on-target off-tumor and/or graft-versus-host disease.

In conclusion, this study demonstrated the efficacy of SSEA-4 CAR-T cells in an HGSOC pre-clinical model. However, SSEA-4 CAR-T cell therapy produced severe adverse effects, therefore further assessment and/or toxicity management strategies will be required before its further clinical translation. Alternative strategies could involve lowering the CAR-T cell dose or scFv affinity, equipping CAR-T cells with combinatorial antigen recognition [43, 44] such as SSEA-4 and mesothelin [45], and/or sigmoidal antigen-density threshold [46] using low-affinity synthetic Notch receptors to control the expression of the high-affinity anti-SSEA-4 CAR.

## Abbreviations

CAR-T: Chimeric antigen receptor T-cell
scFv: Single-chain variable fragment
TME: Tumor microenvironment
ECM: Extracellular matrix
TAA: Tumor-associated antigen
CSC: Cancer stem-like cell
ADCC: Antibody-dependent cytotoxicity
SSEA-4: Stage-specific embryonic antigen-4
GSL: Glycosphingolipid
SC: Stem cell
HGSOC: High-grade serous ovarian carcinomas
NSG: NOD scid gamma
BLI: Bioluminescence imaging

## Ethics approval

Samples from patients with metastatic HGSC were obtained at The Karolinska University Hospital under protocols approved by The Swedish Ethical Review Agency (Etikprövningsmyndigheten; 2016/119731/1, 2016/2060-32). All methods for the use of human samples were carried out in accordance with relevant guidelines and regulations. Informed consent was obtained from all the patients. For gastric and pancreatic cancer samples, the study has the approval of the Helsinki University Hospital Ethics Committee (document number HUS 226/E6/06, additional petition TMK02 § 66/2013 and HUS/1223/2021). Permission to use archive material for this retrospective study without individual informed consent from the patients has been granted by the Finnish National Supervisory Authority for Welfare and Health (document number 1004/06.01.03.01/2012) and the Finnish Medicines Agency Fimea (document number FIMEA/2021/006901).

## Authors’ contributions

**H.J.M**. and **M.H**. Conceptualization, formal analysis, validation, investigation, methodology, writing-original draft, writing-review and editing. **E.E**. Formal analysis, validation, investigation, methodology, writing-review and editing. **K.K**., **J.W**., **L.M-G**. and **A.R**. Resources, investigation, methodology, writing-review and editing. **T.R.J**. and **J.J**. Investigation, methodology. **P.L** and **K.L**. Resources, writing-review and editing. **H.S**., **P.P**., **J.W.C**., **S.S**. and **C.H**. Resources, writing-review and editing. **P.M.O**. and **S.L**. Conceptualization, resources, formal analysis, funding acquisition, validation, writing-original draft, writing-review and editing.

## Consent for publication

All contributors have agreed to publish this manuscript.

## Data availability

The datasets presented in this study are available from the corresponding authors with reasonable requests. This study adheres to ICMJE standards.

## Acknowledgments

We thank the Laboratory Animal Center (LAC), Biomedicum Imaging Unit (BIU), Genome Biology Unit (GBU) at the University of Helsinki and the Digital and Molecular Pathology Unit supported by Helsinki University and Biocenter Finland for digital microscopy services for support in animal care and imaging. We are also extremely grateful to Mari Rissanen, Nadezhda Zinovkina, Anne Aarnio, and Sanna Vainionpää (University of Helsinki) for the valuable technical help. This study was supported by CHO Pharma USA Inc. Additional funding was received from the Finnish Cancer Foundation (P.M.O.), Finnish Red Cross Blood Service Research grant (P.M.O., S.L.) and Sigrid Juselius Foundation (P.M.O., S.L.). E.E. was supported by the Academy of Finland Postdoctoral Fellowships (no. 324487 (E.E.). These additional funders had no role in the study design, data collection, interpretation, or decision to submit the work for publication.

## Figure legends

**Supplementary Figure S1. SSEA-4 staining specificity and sensitivity to xylene. A**. SSEA-4 immunohistochemistry in paraffin-embedded gastric cancer tumor microarrays. **B**. Upper panel sections were treated with standard xylene/ethanol series used in paraffin embedded section treatment, whereas lower panel samples were untreated. **C**. Consecutive sections of fresh frozen pancreatic cancer patient samples were stained with anti-SSEA-4 antibody. The staining was blocked with SSEA-4 hexaose (middle panel) or SSEA-3 (right panel). **D**. Representative immunofluorescence micrographs of SSEA-4 (red) in OVCAR-4 cells (high expression) and Caov-3 cells (no expression). Nucleus (blue; Hoechst 33342). N=3.

**Supplementary Figure S2. Short-term *in vivo* study, dose-response, and weight monitoring. A**. Treatment groups and used T cell doses. **B**. Weight monitoring in the indicated mice groups. CAR-T 3M treated mice showed a trend of mild weight loss. (M=1×10^6^).

**Supplementary Figure S3. IFN**γ **measurement and organ BLI of the short-term *in vivo* study**.

**A**. IFNγ quantification by ELISA in post-mortem mouse blood plasma. **B**. Representative BLI images of isolated organs from mice treated with PBS, NT-T, and CAR-T cells at the indicated T cell doses after 32 days. **C**. Organ BLI quantification. N=3 for control. N=5 for all groups, except N=2 for NT-T 1M, N=3 for CAR-T 1M, and N=4 NT-T 2M. Mean ± SD. One-way ANOVA with Tukey’s multiple comparison tests. (**A**) *\*\*\*p=0*.*0005, ****p=0*.*0001*. (**C**) \**p=0*.*04, **p*≤*0*.*001* to PBS group. Not significant, n.s., differences were attributed to *p*≥*0*.*05*. (M=1×10^6^).

**Supplementary Figure S4. Anti-tumor response and survival in the long-term *in vivo* study. A**. Treatment groups and used T cell doses. **B**. Representative BLI images of mice treated with PBS, 1M, and 2M NT or CAR T cells doses at the indicated time points (days; d). (M=1×10^6^).

**Supplementary Figure S5. Weight monitoring during the long-term *in vivo* study**. Mice treated with 2M CAR-T dose showed pronounced weight loss and health deterioration at earlier time points than mice treated with 1M CAR-T dose. PBS, tumor-free CAR/NT-T 1M, and NT-T groups did not show weight loss and remained healthy during the duration of the experiment. Unless otherwise indicated, mice groups are tumor-bearing. (M=1×10^6^).

**Supplementary Figure S6. Expression of SSEA-4 in normal mouse tissues**. Immunofluorescence detection of SSEA-4 (green) in the indicated tissues of healthy 10 weeks old (C57BL/6) mouse using anti-SSEA-4 antibody.

**Supplementary Figure S7. Histopathological analysis of mice after long-term *in vivo* study treated with 2M dose**. H&E staining of mouse tissues from CAR/NT-T treated mice shows prominent lymphocyte infiltrates in the CAR-T treated mice lungs. (M=1×10^6^).

**Supplementary Figure S8. T cell detection in mouse tissues after long term survival study**. Indicated tissues were stained with HOECHST 33342 for nuclei (blue), anti-EGFR to detect CAR-T cells (middle panel; green) and anti-CD3 for T cells (right panel, white). Left panel shows merged image showing all three channels. Arrows highlight CAR-T and CD3 positive cells, whereas arrowheads highlight erythrocytes with intense background staining.

## References

[1] A.F. Ochsenbein, Principles of tumor immunosurveillance and implications for immunotherapy, Cancer Gene Therapy, 9 (2002) 1043–1055.

[2] U. Greenbaum, K.M. Mahadeo, P. Kebriaei, E.J. Shpall, N.Y. Saini, Chimeric Antigen Receptor T-Cells in B-Acute Lymphoblastic Leukemia: State of the Art and Future Directions, 10 (2020).

[3] A.D. Waldman, J.M. Fritz, M.J. Lenardo, A guide to cancer immunotherapy: from T cell basic science to clinical practice, Nature Reviews Immunology, 20 (2020) 651–668.

[4] Z. Walsh, Y. Yang, M.E. Kohler, Immunobiology of chimeric antigen receptor T cells and novel designs, 290 (2019) 100–113.

[5] N.M. Anderson, M.C. Simon, The tumor microenvironment, Current Biology, 30 (2020) R921–R925.

[6] S. Rafiq, C.S. Hackett, R.J. Brentjens, Engineering strategies to overcome the current roadblocks in CAR T cell therapy, Nature Reviews Clinical Oncology, 17 (2020) 147–167.

[7] A. Rodriguez-Garcia, A. Palazon, E. Noguera-Ortega, D.J. Powell, S. Guedan, CAR-T Cells Hit the Tumor Microenvironment: Strategies to Overcome Tumor Escape, 11 (2020).

[8] J. Wagner, E. Wickman, C. DeRenzo, S. Gottschalk, CAR T Cell Therapy for Solid Tumors: Bright Future or Dark Reality?, Mol Ther, 28 (2020) 2320–2339.

[9] K. Newick, S. O’Brien, E. Moon, S.M. Albelda, CAR T Cell Therapy for Solid Tumors, 68 (2017) 139–152.

[10] T.N. Yamamoto, R.J. Kishton, N.P. Restifo, Developing neoantigen-targeted T cell–based treatments for solid tumors, Nature Medicine, 25 (2019) 1488–1499.

[11] R.C. Sterner, R.M. Sterner, CAR-T cell therapy: current limitations and potential strategies, Blood Cancer Journal, 11 (2021) 69.

[12] L.J.B. Brandt, M.B. Barnkob, Y.S. Michaels, J. Heiselberg, T. Barington, Emerging Approaches for Regulation and Control of CAR T Cells: A Mini Review, 11 (2020).

[13] J.N. Brudno, J.N. Kochenderfer, Toxicities of chimeric antigen receptor T cells: recognition and management, Blood, 127 (2016) 3321–3330.

[14] N. Dasyam, P. George, R. Weinkove, Chimeric antigen receptor T-cell therapies: Optimising the dose, Br J Clin Pharmacol, 86 (2020) 1678–1689.

[15] M. Addeo, G. Di Paola, H.K. Verma, S. Laurino, S. Russi, P. Zoppoli, G. Falco, P. Mazzone, Gastric Cancer Stem Cells: A Glimpse on Metabolic Reprogramming, 11 (2021).

[16] N. Terraneo, F. Jacob, A. Dubrovska, J. Grünberg, Novel Therapeutic Strategies for Ovarian Cancer Stem Cells, 10 (2020).

[17] G. Askan, I.H. Sahin, J.F. Chou, A. Yavas, M. Capanu, C.A. Iacobuzio-Donahue, O. Basturk, E.M. O’Reilly, Pancreatic cancer stem cells may define tumor stroma characteristics and recurrence patterns in pancreatic ductal adenocarcinoma, BMC Cancer, 21 (2021) 385.

[18] R.Y. Alhabbab, Targeting Cancer Stem Cells by Genetically Engineered Chimeric Antigen Receptor T Cells, Front Genet, 11 (2020) 312.

[19] B.C. Prager, Q. Xie, S. Bao, J.N. Rich, Cancer Stem Cells: The Architects of the Tumor Ecosystem, Cell Stem Cell, 24 (2019) 41–53.

[20] D. Russo, L. Capolupo, J.S. Loomba, L. Sticco, G. D’Angelo, Glycosphingolipid metabolism in cell fate specification, Journal of Cell Science, 131 (2018).

[21] J. Munkley, D.J. Elliott, Hallmarks of glycosylation in cancer, 7 (2016).

[22] K. Sivasubramaniyan, A. Harichandan, K. Schilbach, A.F. Mack, J. Bedke, A. Stenzl, L. Kanz, G. Niederfellner, H.-J. Bühring, Expression of stage-specific embryonic antigen-4 (SSEA-4) defines spontaneous loss of epithelial phenotype in human solid tumor cells, Glycobiology, 25 (2015) 902–917.

[23] M.-Y. Ho, A.L. Yu, J. Yu, Glycosphingolipid dynamics in human embryonic stem cell and cancer: their characterization and biomedical implications, Glycoconjugate Journal, 34 (2017) 765–777.

[24] S.C. Parte, S.K. Batra, S.S. Kakar, Characterization of stem cell and cancer stem cell populations in ovary and ovarian tumors, J Ovarian Res, 11 (2018) 69–69.

[25] C. Soliman, J.X. Chua, M. Vankemmelbeke, R.S. McIntosh, A.J. Guy, I. Spendlove, L.G. Durrant, P.A. Ramsland, The terminal sialic acid of stage-specific embryonic antigen-4 has a crucial role in binding to a cancer-targeting antibody, Journal of Biological Chemistry, 295 (2020) 1009–1020.

[26] Y.-W. Lou, P.-Y. Wang, S.-C. Yeh, P.-K. Chuang, S.-T. Li, C.-Y. Wu, K.-H. Khoo, M. Hsiao, T.-L. Hsu, C.-H. Wong, Stage-specific embryonic antigen-4 as a potential therapeutic target in glioblastoma multiforme and other cancers, Proceedings of the National Academy of Sciences, 111 (2014) 2482.

[27] O. Firuzi, P.P. Che, B. El Hassouni, M. Buijs, S. Coppola, M. Löhr, N. Funel, R. Heuchel, I. Carnevale, T. Schmidt, G. Mantini, A. Avan, L. Saso, G.J. Peters, E. Giovannetti, Role of c-MET Inhibitors in Overcoming Drug Resistance in Spheroid Models of Primary Human Pancreatic Cancer and Stellate Cells, 11 (2019) 638.

[28] D.S. Sigal, D.J. Hermel, P. Hsu, T. Pearce, The role of Globo H and SSEA-4 in the development and progression of cancer, and their potential as therapeutic targets, Future Oncol, 18 (2022) 117–134.

[29] C.W. Lin, Y.J. Wang, T.Y. Lai, T.L. Hsu, S.Y. Han, H.C. Wu, C.N. Shen, V. Dang, M.W. Chen, L.B. Chen, C.H. Wong, Homogeneous antibody and CAR-T cells with improved effector functions targeting SSEA-4 glycan on pancreatic cancer, Proc Natl Acad Sci U S A, 118 (2021).

[30] X. Wang, W.-C. Chang, C.W. Wong, D. Colcher, M. Sherman, J.R. Ostberg, S.J. Forman, S.R. Riddell, M.C. Jensen, A transgene-encoded cell surface polypeptide for selection, in vivo tracking, and ablation of engineered cells, Blood, 118 (2011) 1255–1263.

[31] A. Magalhães, H.O. Duarte, C.A. Reis, Aberrant Glycosylation in Cancer: A Novel Molecular Mechanism Controlling Metastasis, Cancer Cell, 31 (2017) 733–735.

[32] R. Zhou, M. Yazdanifar, L.D. Roy, L.M. Whilding, A. Gavrill, J. Maher, P. Mukherjee, CAR T Cells Targeting the Tumor MUC1 Glycoprotein Reduce Triple-Negative Breast Cancer Growth, Frontiers in Immunology, 10 (2019).

[33] M. Prapa, C. Chiavelli, G. Golinelli, G. Grisendi, M. Bestagno, R. Di Tinco, M. Dall’Ora, G. Neri, O. Candini, C. Spano, T. Petrachi, L. Bertoni, G. Carnevale, G. Pugliese, R. Depenni, A. Feletti, C. Iaccarino, G. Pavesi, M. Dominici, GD2 CAR T cells against human glioblastoma, npj Precision Oncology, 5 (2021) 93.

[34] Y.W. Lou, P.Y. Wang, S.C. Yeh, P.K. Chuang, S.T. Li, C.Y. Wu, K.H. Khoo, M. Hsiao, T.L. Hsu, C.H. Wong, Stage-specific embryonic antigen-4 as a potential therapeutic target in glioblastoma multiforme and other cancers, Proc Natl Acad Sci U S A, 111 (2014) 2482–2487.

[35] J. Masoumi, A. Jafarzadeh, J. Abdolalizadeh, H. Khan, J. Philippe, H. Mirzaei, H.R. Mirzaei, Cancer stem cell-targeted chimeric antigen receptor (CAR)-T cell therapy: Challenges and prospects, Acta Pharmaceutica Sinica B, 11 (2021) 1721–1739.

[36] J.W.Y. Wu, S. Dand, L. Doig, A.T. Papenfuss, C.L. Scott, G. Ho, J.D. Ooi, T-Cell Receptor Therapy in the Treatment of Ovarian Cancer: A Mini Review, Frontiers in immunology, 12 (2021) 672502–672502.

[37] X. Zhou, L. Rasche, K.M. Kortüm, S. Danhof, M. Hudecek, H. Einsele, Toxicities of Chimeric Antigen Receptor T Cell Therapy in Multiple Myeloma: An Overview of Experience From Clinical Trials, Pathophysiology, and Management Strategies, Frontiers in Immunology, 11 (2020).

[38] P.H. Mehta, S. Fiorenza, R.M. Koldej, A. Jaworowski, D.S. Ritchie, K.M. Quinn, T Cell Fitness and Autologous CAR T Cell Therapy in Haematologic Malignancy, Frontiers in Immunology, 12 (2021).

[39] D.J. Green, M. Pont, B.D. Sather, A.J. Cowan, C.J. Turtle, B.G. Till, A.M. Nagengast, E.N. Libby, III, P.S. Becker, D.G. Coffey, S.A. Tuazon, B. Wood, M. Blake, M. Works, B.S. Thompson, T. Gooley, F.R. Appelbaum, D.G. Maloney, S.R. Riddell, Fully Human Bcma Targeted Chimeric Antigen Receptor T Cells Administered in a Defined Composition Demonstrate Potency at Low Doses in Advanced Stage High Risk Multiple Myeloma, Blood, 132 (2018) 1011–1011.

[40] S. Jiang, J. Jin, S. Hao, M. Yang, L. Chen, H. Ruan, J. Xiao, W. Wang, Z. Li, K. Yu, Low Dose of Human scFv-Derived BCMA-Targeted CAR-T Cells Achieved Fast Response and High Complete Remission in Patients with Relapsed/Refractory Multiple Myeloma, Blood, 132 (2018) 960–960.

[41] X. Cui, R. Liu, L. Duan, D. Cao, Q. Zhang, A. Zhang, CAR-T therapy: Prospects in targeting cancer stem cells, Journal of Cellular and Molecular Medicine, 25 (2021) 9891–9904.

[42] M.L. Disis, H. Bernhard, E.M. Jaffee, Use of tumour-responsive T cells as cancer treatment, Lancet, 373 (2009) 673–683.

[43] C.C. Kloss, M. Condomines, M. Cartellieri, M. Bachmann, M. Sadelain, Combinatorial antigen recognition with balanced signaling promotes selective tumor eradication by engineered T cells, Nat Biotechnol, 31 (2013) 71–75.

[44] J.H. Choe, P.B. Watchmaker, M.S. Simic, R.D. Gilbert, A.W. Li, N.A. Krasnow, K.M. Downey, W. Yu, D.A. Carrera, A. Celli, J. Cho, J.D. Briones, J.M. Duecker, Y.E. Goretsky, R. Dannenfelser, L. Cardarelli, O. Troyanskaya, S.S. Sidhu, K.T. Roybal, H. Okada, W.A. Lim, SynNotch-CAR T cells overcome challenges of specificity, heterogeneity, and persistence in treating glioblastoma, Sci Transl Med, 13 (2021).

[45] E. Schoutrop, I. El-Serafi, T. Poiret, Y. Zhao, O. Gultekin, R. He, L. Moyano-Galceran, J.W. Carlson, K. Lehti, M. Hassan, I. Magalhaes, J. Mattsson, Mesothelin-Specific CAR T Cells Target Ovarian Cancer, Cancer Res, 81 (2021) 3022–3035.

[46] A. Hernandez-Lopez Rogelio, W. Yu, A. Cabral Katelyn, A. Creasey Olivia, P. Lopez Pazmino Maria del, Y. Tonai, A. De Guzman, A. Mäkelä, K. Saksela, J. Gartner Zev, A. Lim Wendell, T cell circuits that sense antigen density with an ultrasensitive threshold, Science, 371 (2021) 1166–1171.

