## Supplementary material for "Efficacy and safety of glycosphingolipid SSEA-4 targeting CAR-T cells in an ovarian carcinoma model": Figure Supplementary S1.

Supplementary Figure S1.

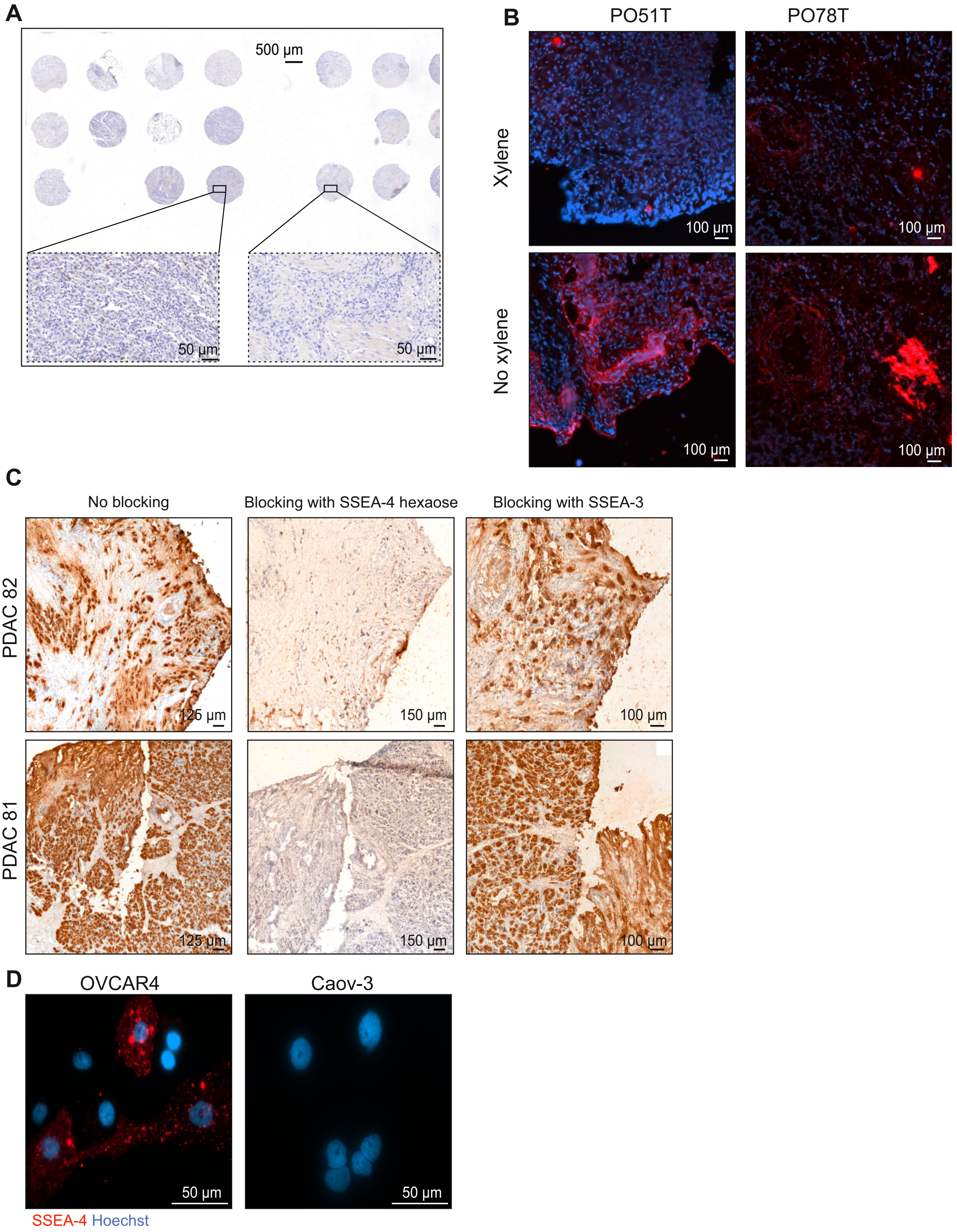

Supplementary Figure S2.

**A**

| Group | Mouse n | OVCAR4 cells | Treatment | Transduction | T cell dose | CAR-T cells | NT-T cells | Total T cells |
|---|---|---|---|---|---|---|---|---|
| 1 | 5 | 2.0x10$^6$ | PBS | --- | --- | --- | --- | --- |
| 2 | 2 | 2.0x10$^6$ | NT-T | --- | 1M | --- | 2.9x10$^6$ | 2.9x10$^6$ |
| | 3 | 2.0x10$^6$ | CAR-T | 35% | | 1x10$^6$ | 1.9x10$^6$ | 2.9x10$^6$ |
| 3 | 5 | 2.0x10$^6$ | NT-T | --- | 2M | --- | 5.7x10$^6$ | 5.7x10$^6$ |
| | 5 | 2.0x10$^6$ | CAR-T | 35% | | 2x10$^6$ | 3.7x10$^6$ | 5.7x10$^6$ |
| 4 | 5 | 2.0x10$^6$ | NT-T | --- | 3M | --- | 8.6x10$^6$ | 8.6x10$^6$ |
| | 5 | 2.0x10$^6$ | CAR-T | 35% | | 3x10$^6$ | 5.6x10$^6$ | 8.6x10$^6$ |
| 5 | 3 | --- | --- | --- | --- | --- | --- | --- |

**B**

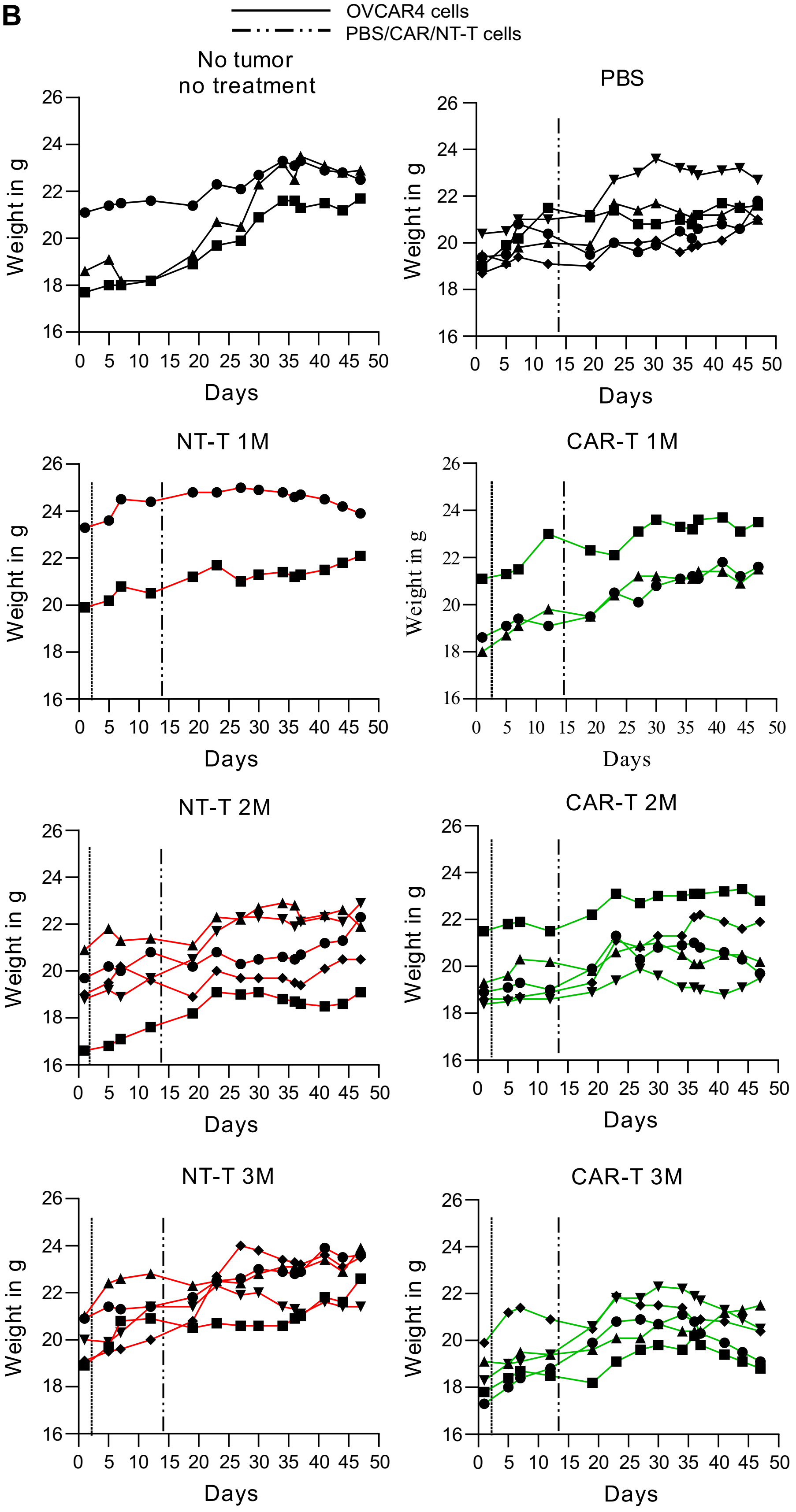

Supplementary Figure S3.

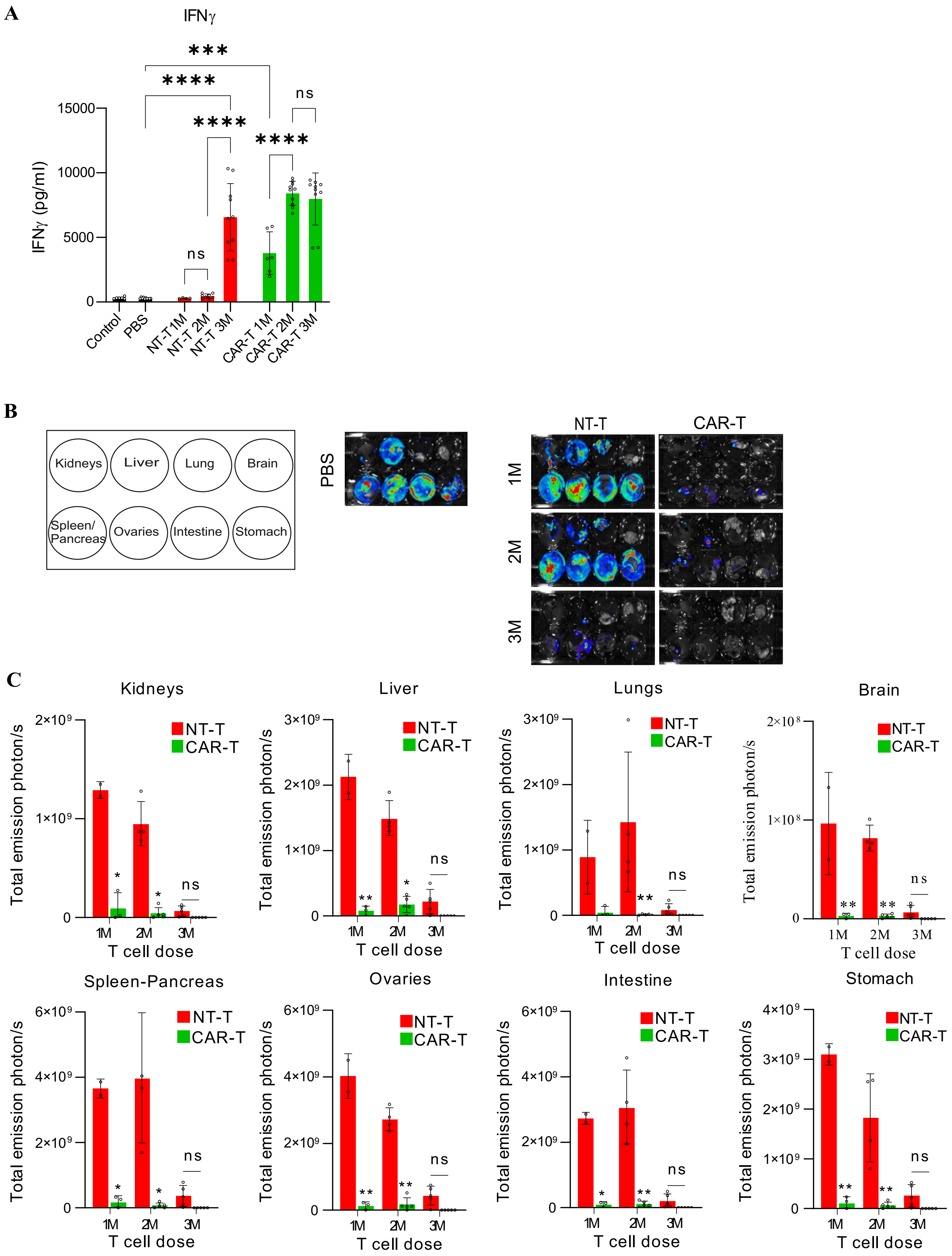

Supplementary Fig. S4.

**A**

| Group | Mouse n | OVCAR4 cells | Treatment | Transduction | T cell dose | CAR-T cells | NT-T cells | Total T cells |
|---|---|---|---|---|---|---|---|---|
| 1 | 10 | $2.0 \times 10^6$ | PBS | --- | --- | --- | --- | --- |
| 2 | 10 | $2.0 \times 10^6$ | NT-T | --- | 1M | --- | $1.1 \times 10^6$ | $1.1 \times 10^6$ |
| | 10 | $2.0 \times 10^6$ | CAR-T | 90% | | $1 \times 10^6$ | $0.1 \times 10^6$ | $1.1 \times 10^6$ |
| 3 | 10 | $2.0 \times 10^6$ | NT-T | --- | 2M | --- | $2.2 \times 10^6$ | $2.2 \times 10^6$ |
| | 10 | $2.0 \times 10^6$ | CAR-T | 90% | | $2 \times 10^6$ | $0.2 \times 10^6$ | $2.2 \times 10^6$ |
| 4 | 2 | --- | NT-T | --- | 1M | --- | $1.1 \times 10^6$ | $1.1 \times 10^6$ |
| 5 | 2 | --- | CAR-T | 90% | | $1 \times 10^6$ | $0.1 \times 10^6$ | $1.1 \times 10^6$ |

**B. 1M**

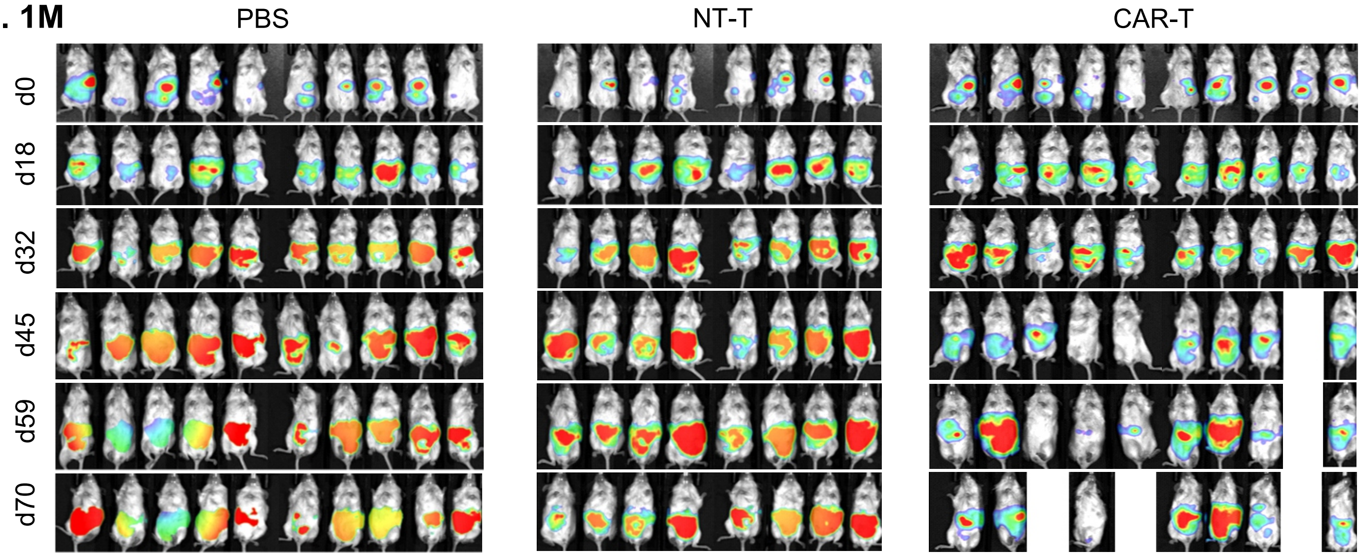


**C. 2M**

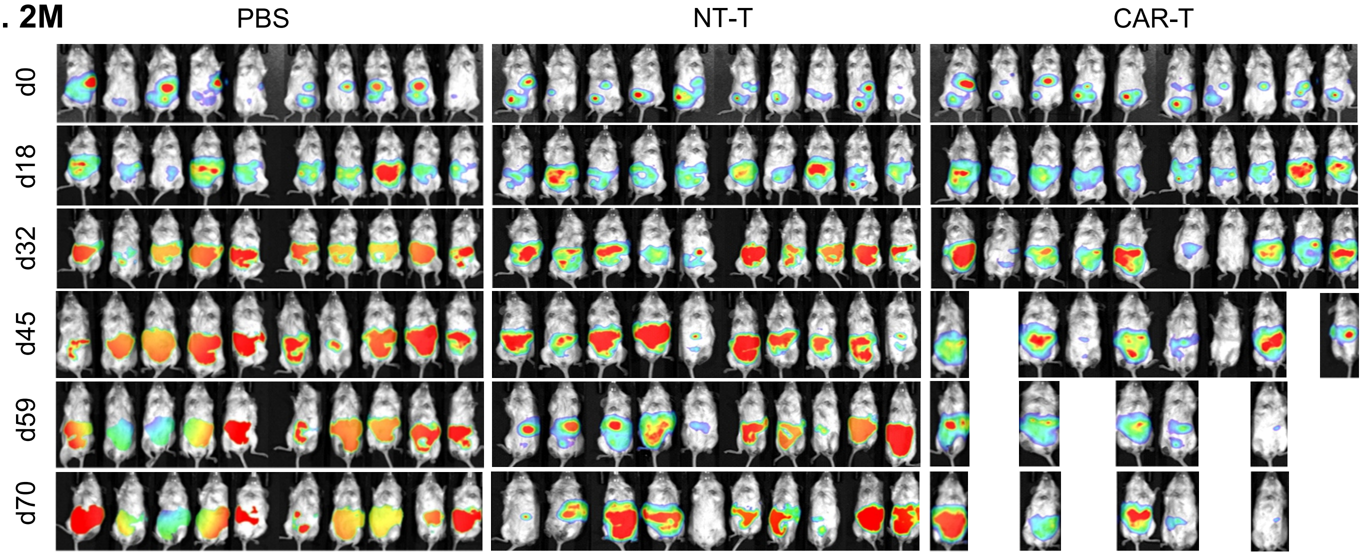

Supplementary Figure S5.

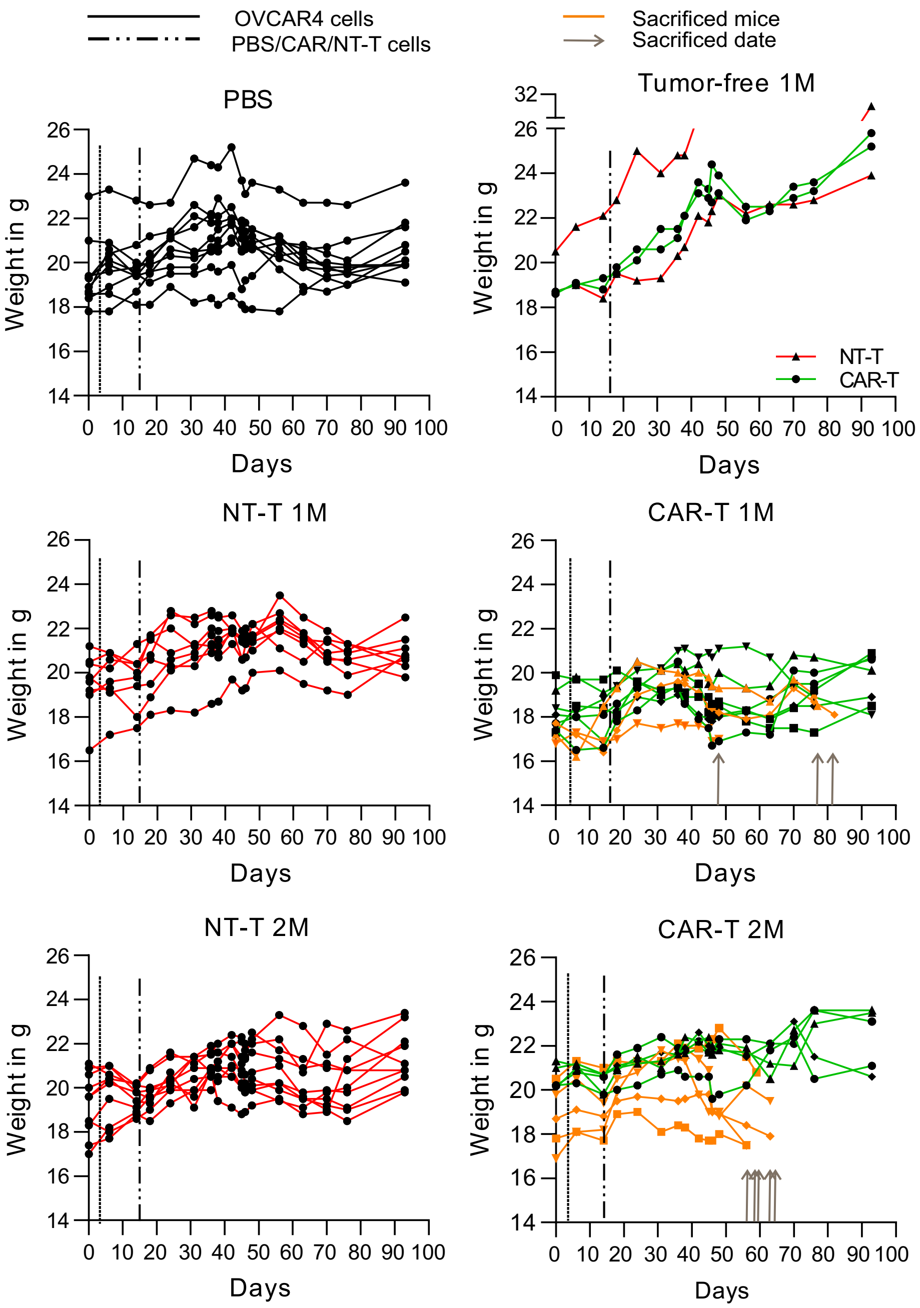

Supplementary Figure S6.

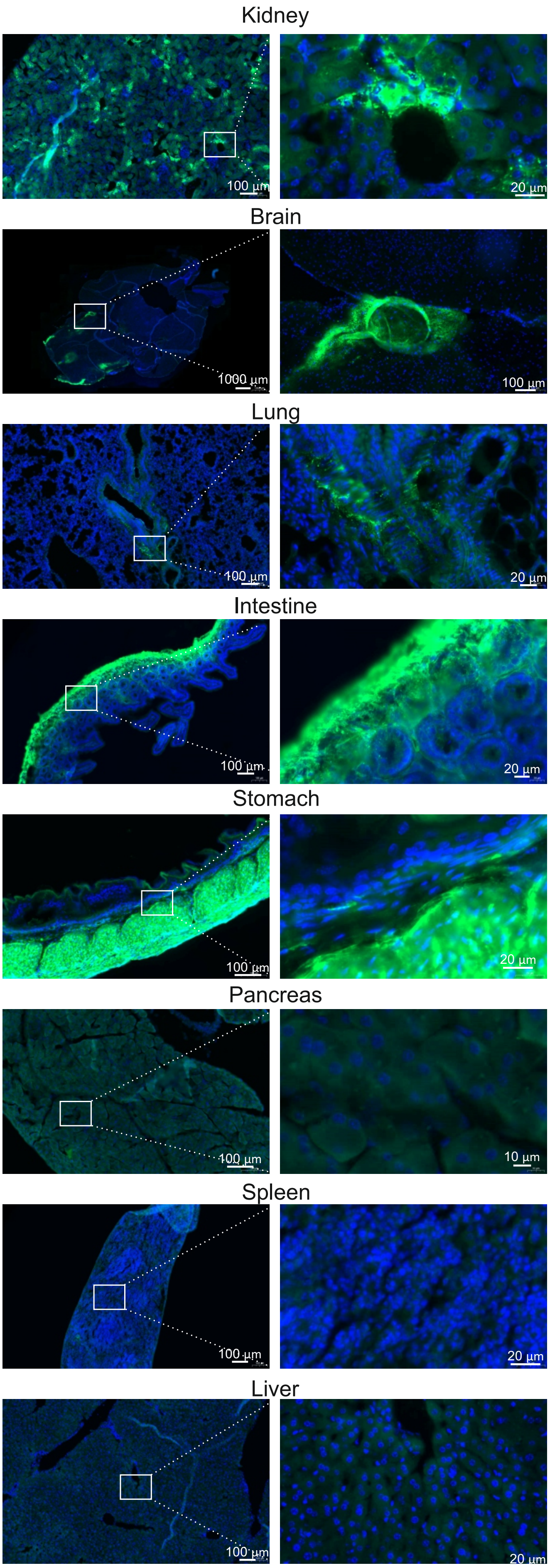

Supplementary Figure S7.

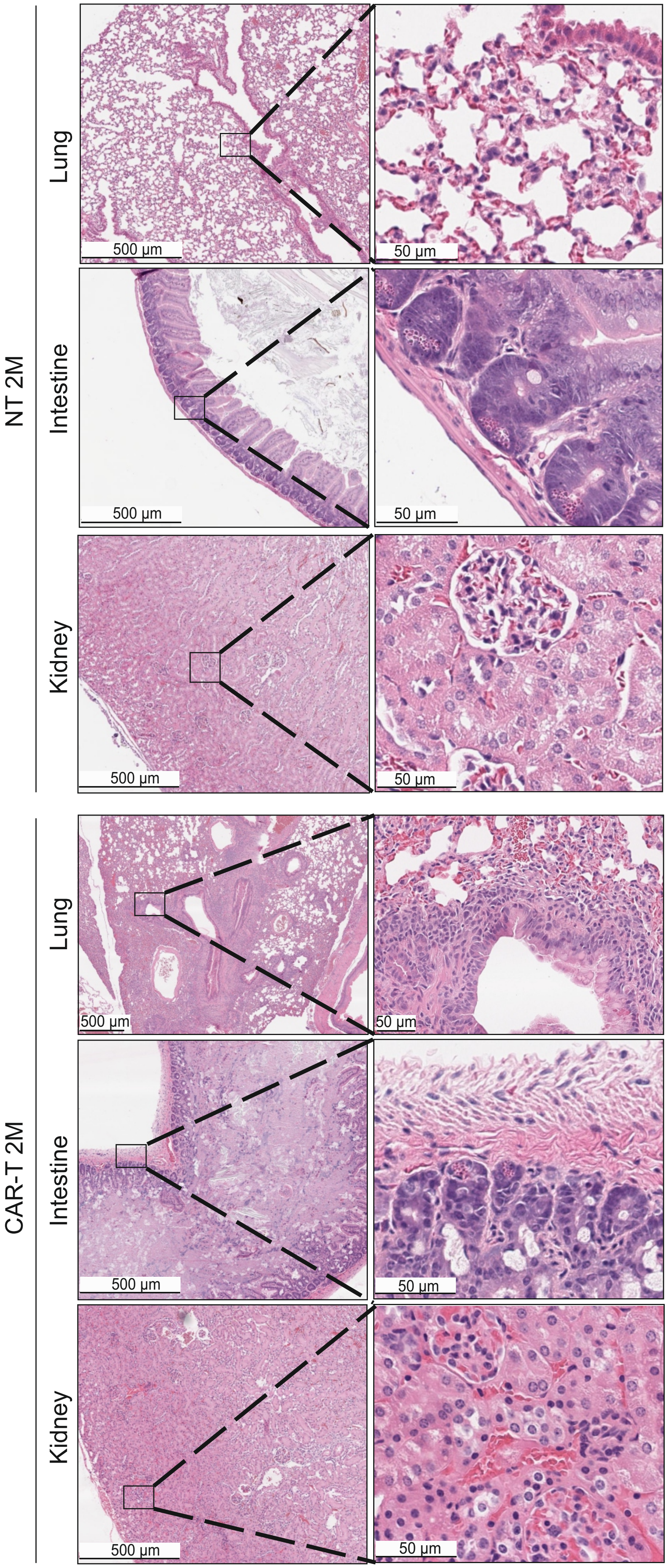

Supplementary Figure S8.

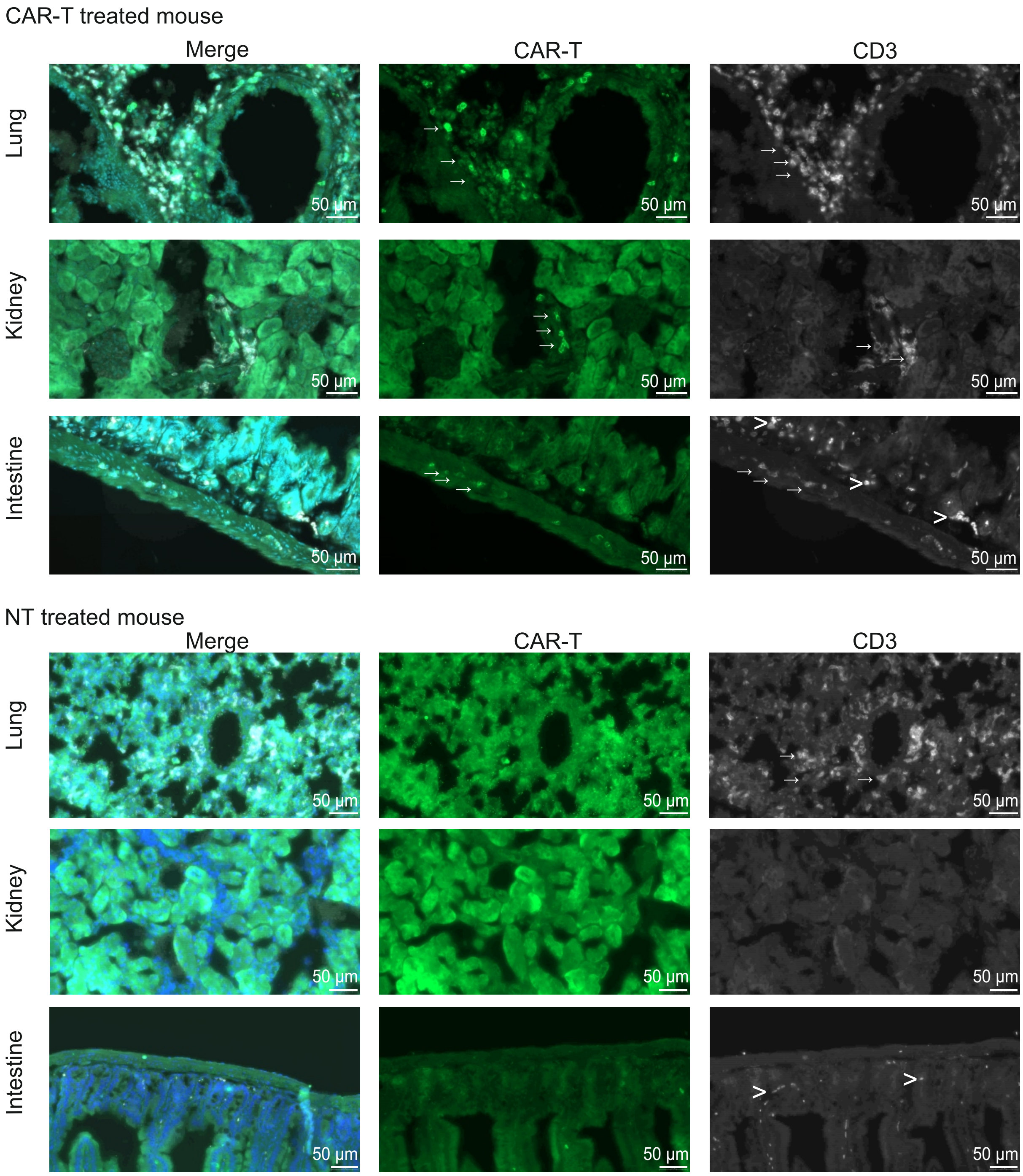
