## Supplementary material for "Efficacy and safety of glycosphingolipid SSEA-4 targeting CAR-T cells in an ovarian carcinoma model": Figure Supplementary S2.

Supplementary Figure S2.

**A**

| Group | Mouse n | OVCAR4 cells | Treatment | Transduction | T cell dose | CAR-T cells | NT-T cells | Total T cells |
|---|---|---|---|---|---|---|---|---|
| 1 | 5 | 2.0x10⁶ | PBS | --- | --- | --- | --- | --- |
| 2 | 2 | 2.0x10⁶ | NT-T | --- | 1M | --- | 2.9x10⁶ | 2.9x10⁶ |
|  | 3 | 2.0x10⁶ | CAR-T | 35% |  | 1x10⁶ | 1.9x10⁶ | 2.9x10⁶ |
| 3 | 5 | 2.0x10⁶ | NT-T | --- | 2M | --- | 5.7x10⁶ | 5.7x10⁶ |
|  | 5 | 2.0x10⁶ | CAR-T | 35% |  | 2x10⁶ | 3.7x10⁶ | 5.7x10⁶ |
| 4 | 5 | 2.0x10⁶ | NT-T | --- | 3M | --- | 8.6x10⁶ | 8.6x10⁶ |
|  | 5 | 2.0x10⁶ | CAR-T | 35% |  | 3x10⁶ | 5.6x10⁶ | 8.6x10⁶ |
| 5 | 3 | --- | --- | --- | --- | --- | --- | --- |

**B**

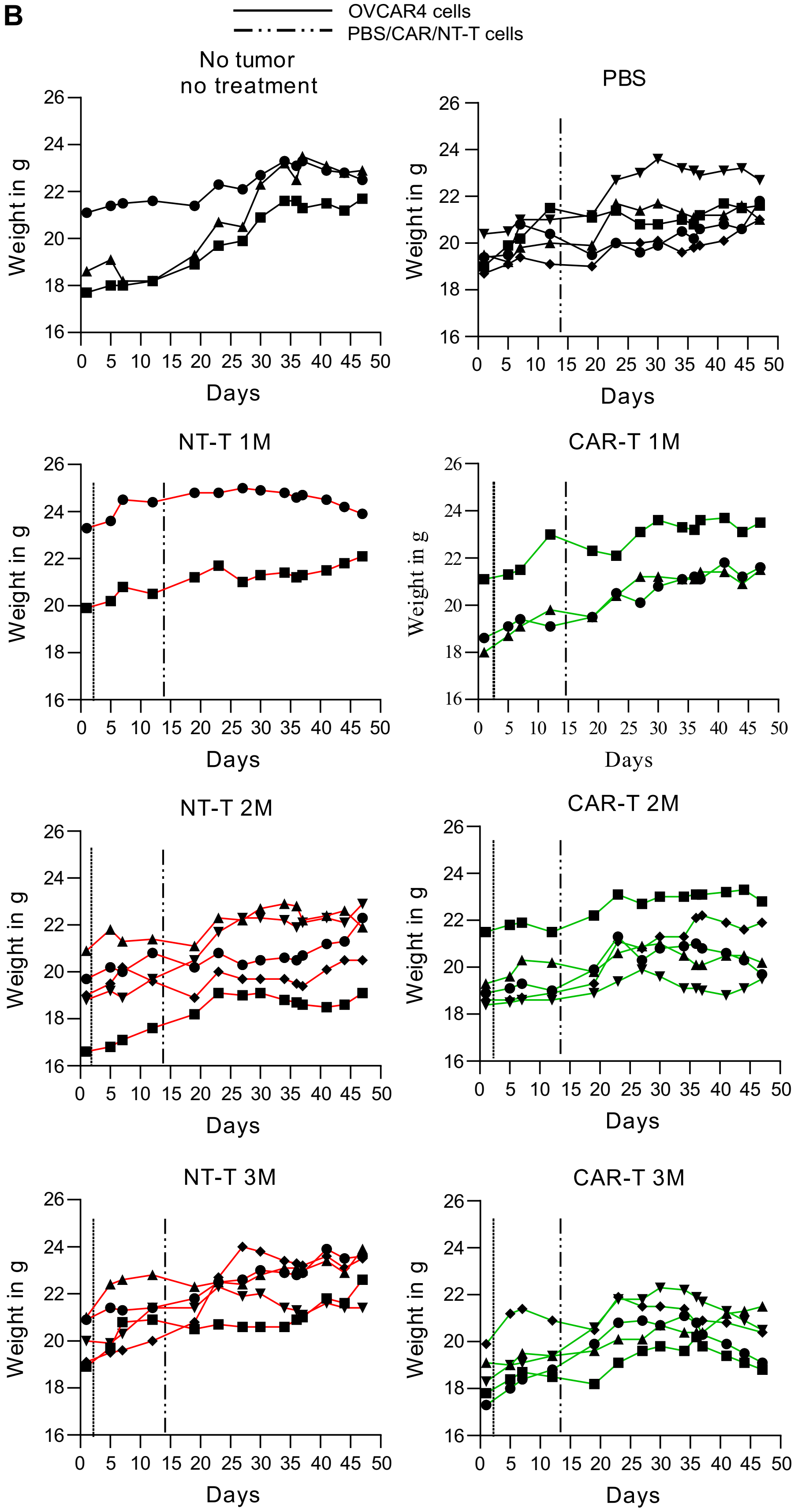
