## Supplementary figures and images for "Efficacy and safety of glycosphingolipid SSEA-4 targeting CAR-T cells in an ovarian carcinoma model"

### Figure Supplementary S3.

Supplementary Figure S3.

A

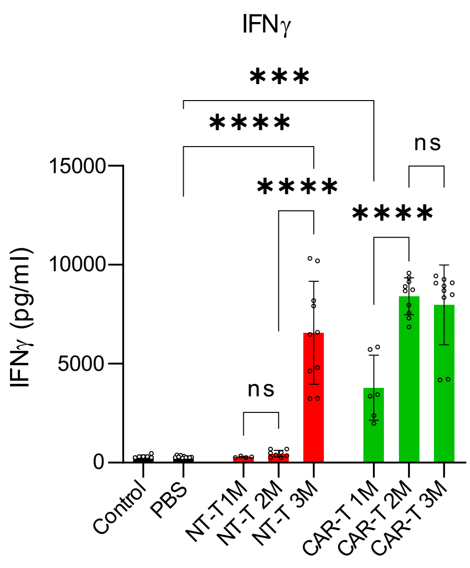


B

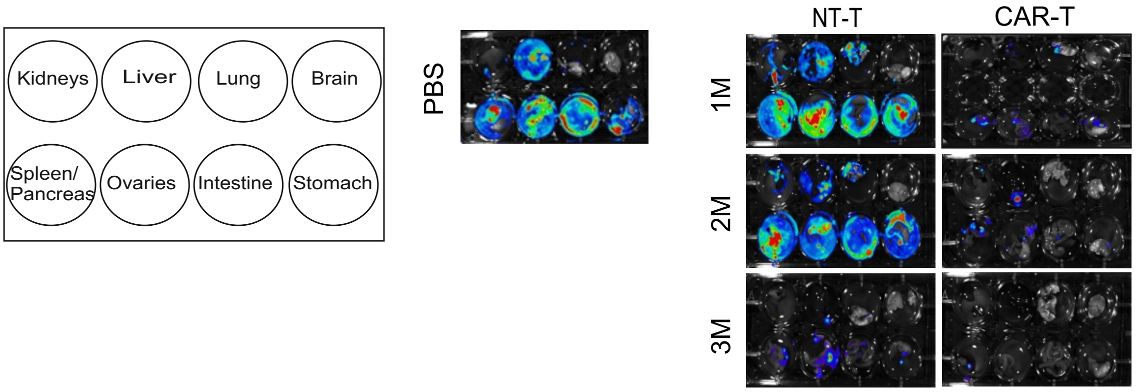


C

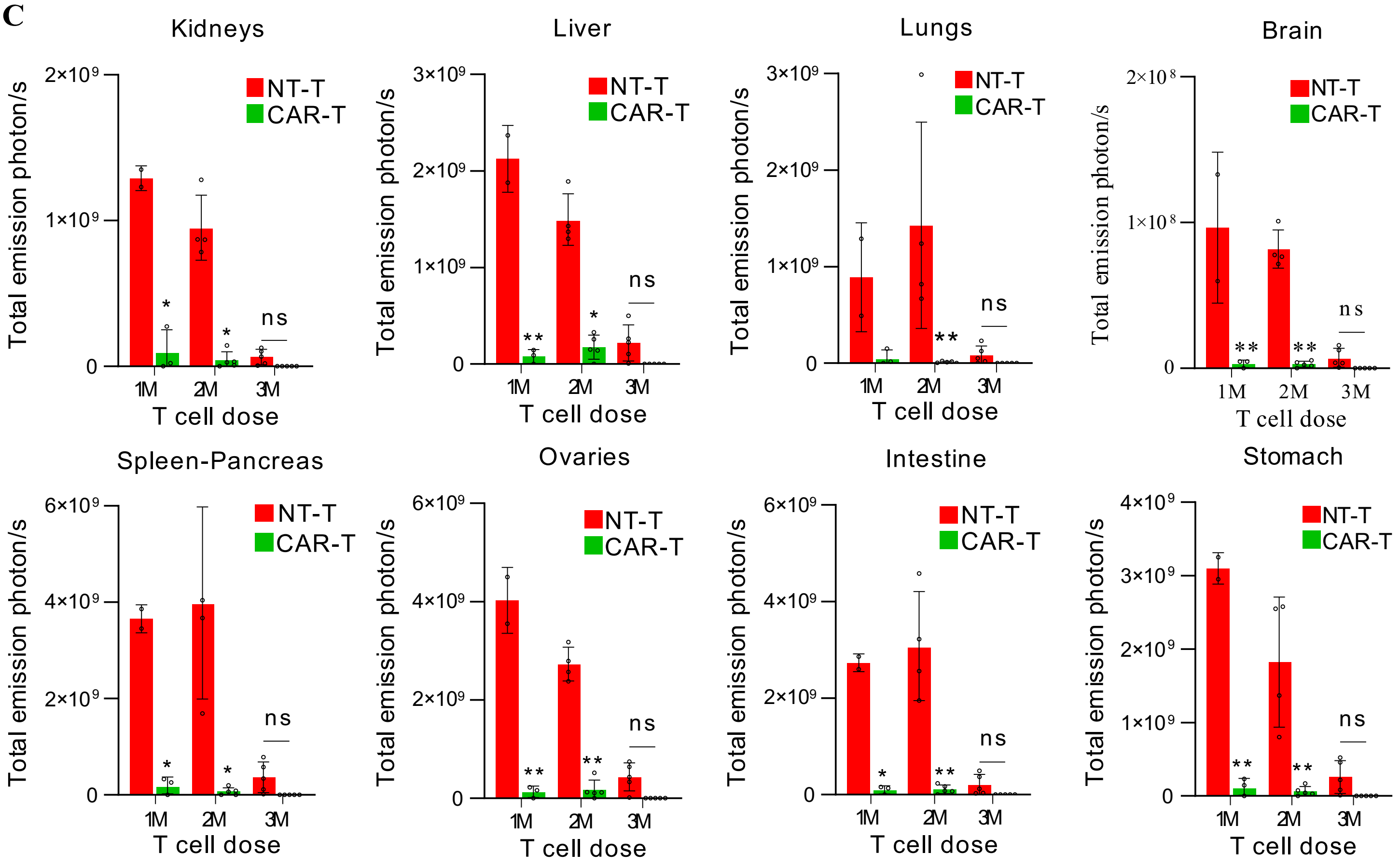

### Figure Supplementary S5.

Supplementary Figure S5.

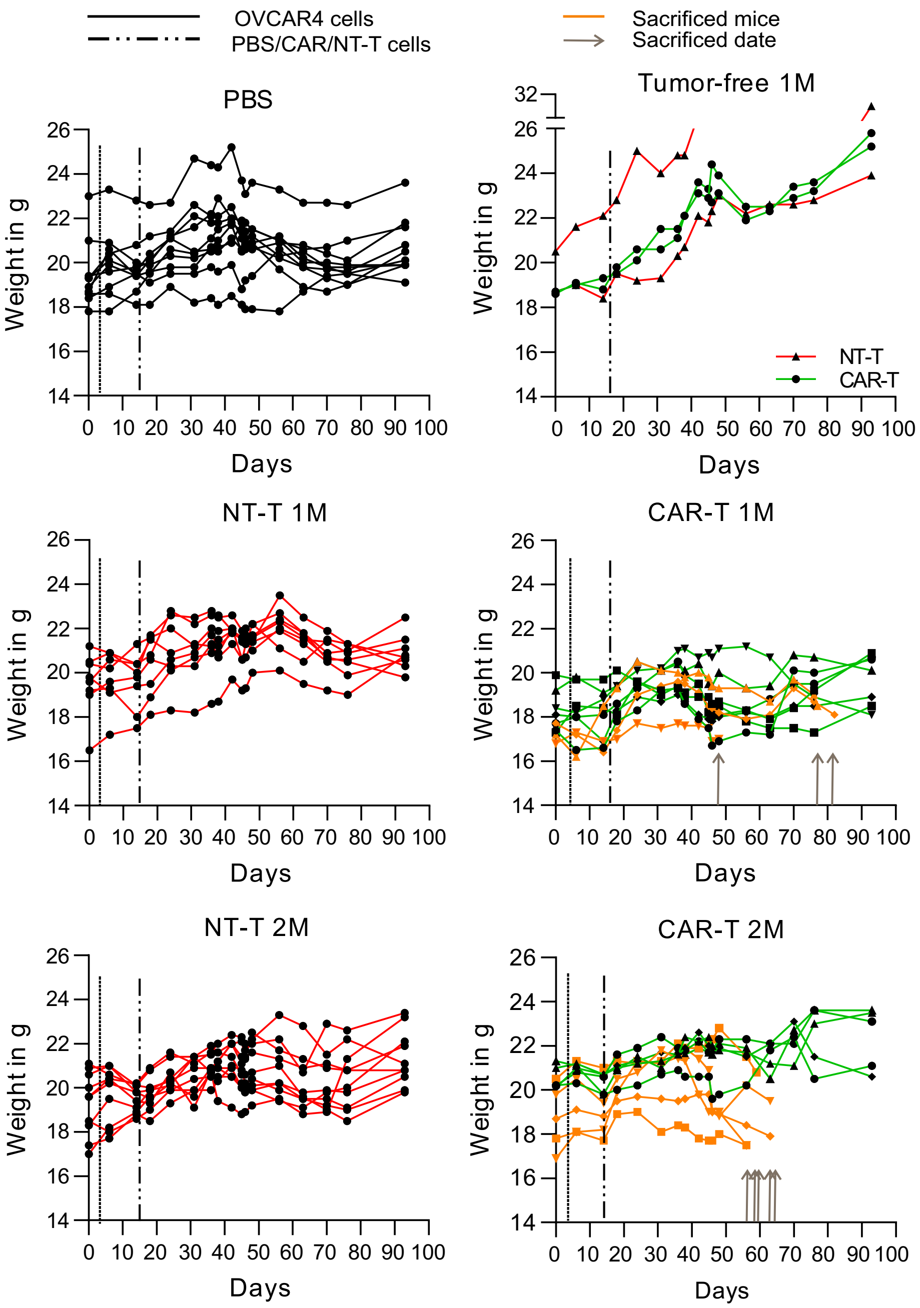

### Figure Supplementary S6.

Supplementary Figure S6.

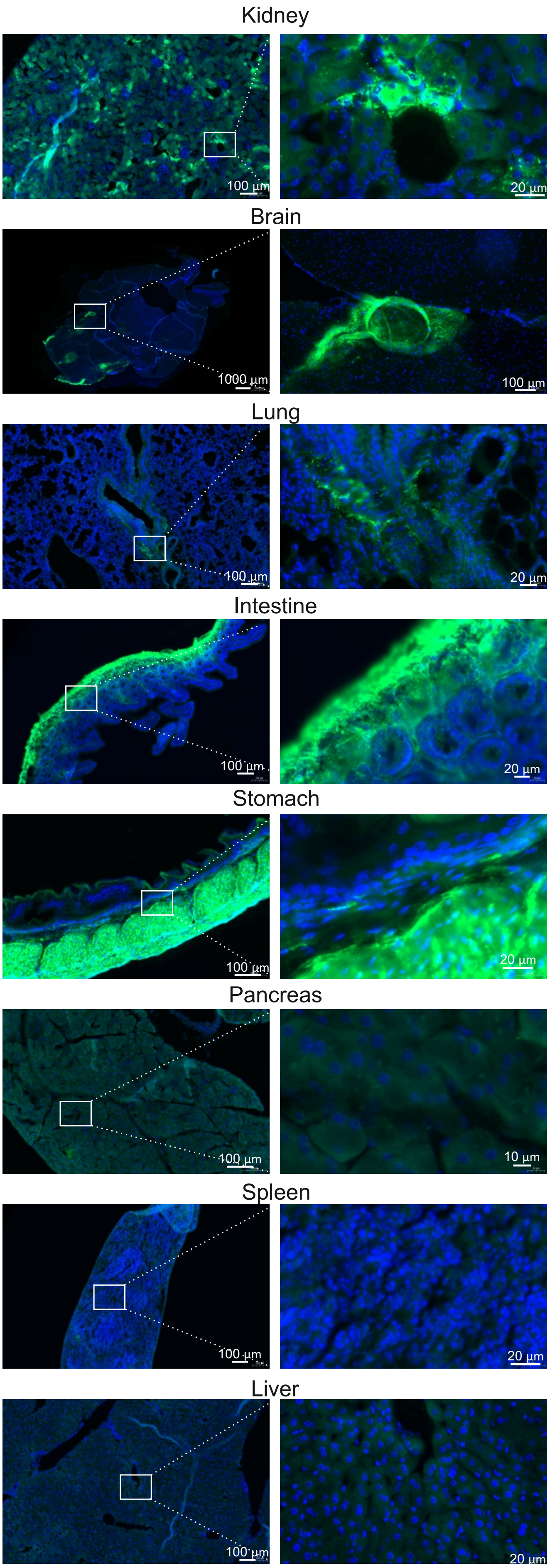

### Figure Supplementary S7.

Supplementary Figure S7.

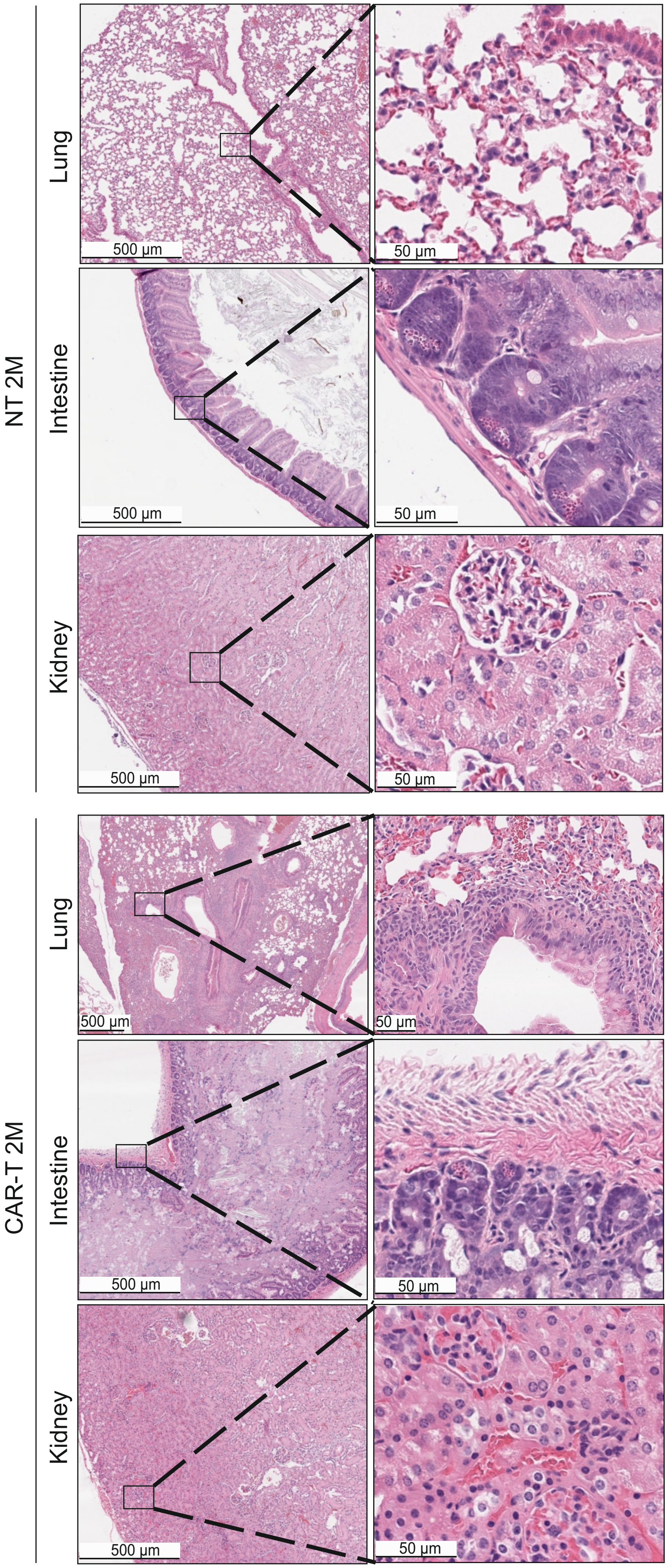

### Figure Supplementary S8.

Supplementary Figure S8.

CAR-T treated mouse

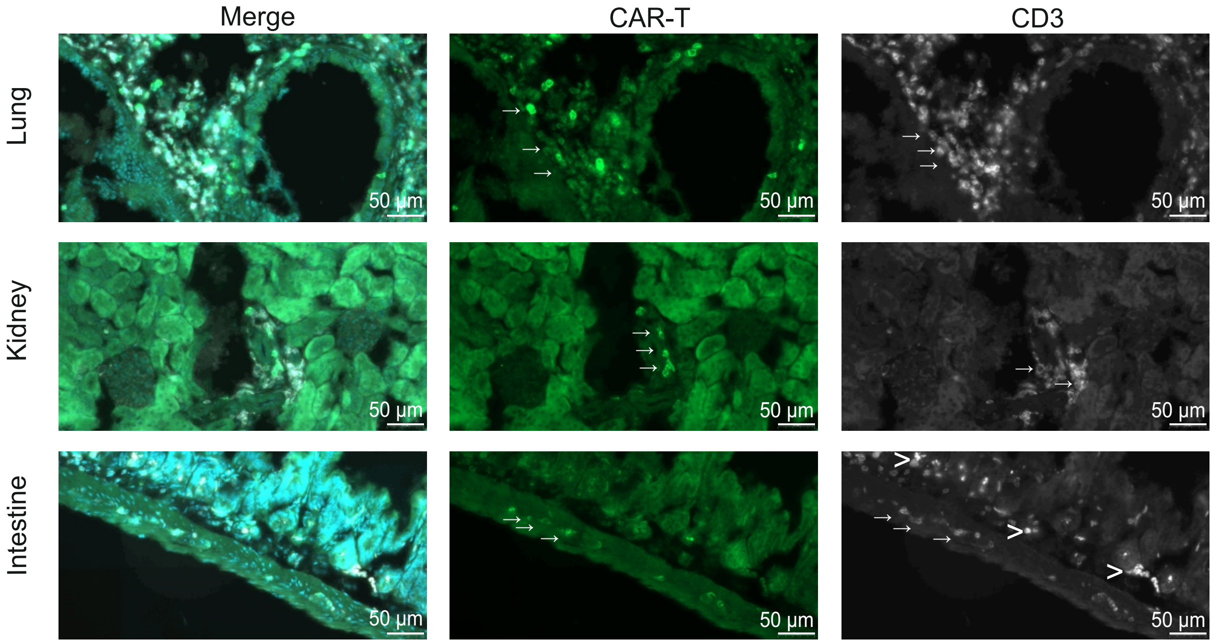


NT treated mouse

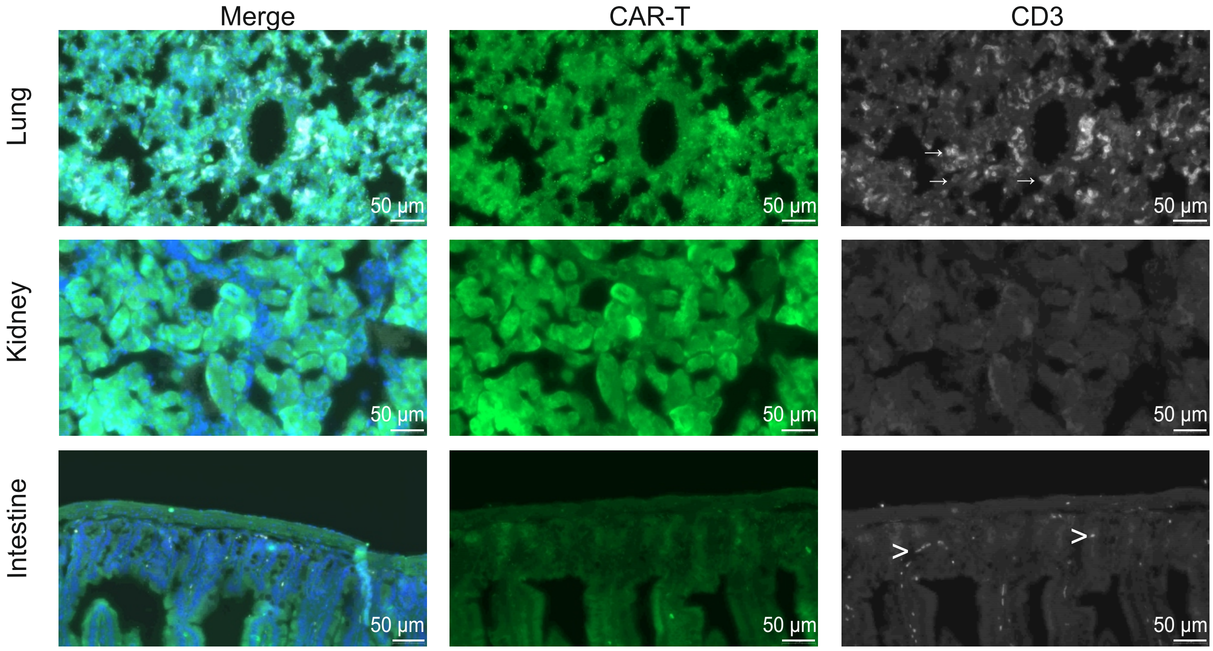
