## Supplementary material for "Efficacy and safety of glycosphingolipid SSEA-4 targeting CAR-T cells in an ovarian carcinoma model": Figure Supplementary S4.

Supplementary Fig. S4.

**A**

| Group | Mouse n | OVCAR4 cells | Treatment | Transduction | T cell dose | CAR-T cells | NT-T cells | Total T cells |
|---|---|---|---|---|---|---|---|---|
| 1 | 10 | $2.0x10^6$ | PBS | --- | --- | --- | --- | --- |
| 2 | 10 | $2.0x10^6$ | NT-T | --- | 1M | --- | $1.1x10^6$ | $1.1x10^6$ |
| | 10 | $2.0x10^6$ | CAR-T | 90% | | $1x10^6$ | $0.1x10^6$ | $1.1x10^6$ |
| 3 | 10 | $2.0x10^6$ | NT-T | --- | 2M | --- | $2.2x10^6$ | $2.2x10^6$ |
| | 10 | $2.0x10^6$ | CAR-T | 90% | | $2x10^6$ | $0.2x10^6$ | $2.2x10^6$ |
| 4 | 2 | --- | NT-T | --- | 1M | --- | $1.1x10^6$ | $1.1x10^6$ |
| 5 | 2 | --- | CAR-T | 90% | | $1x10^6$ | $0.1x10^6$ | $1.1x10^6$ |

**B. 1M**

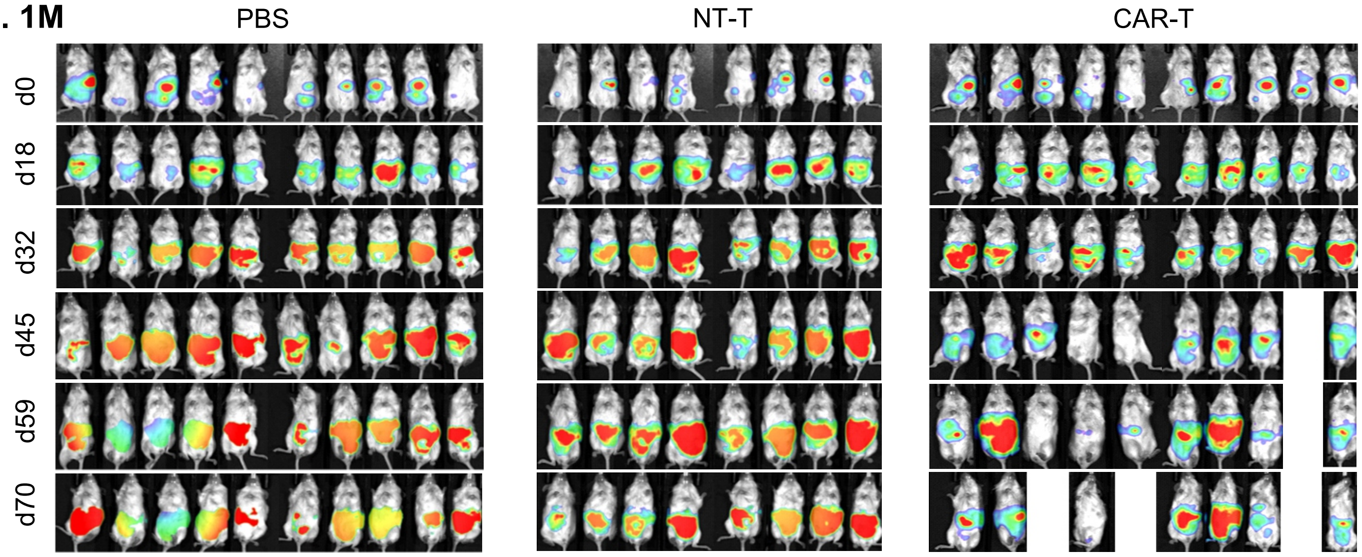


**C. 2M**

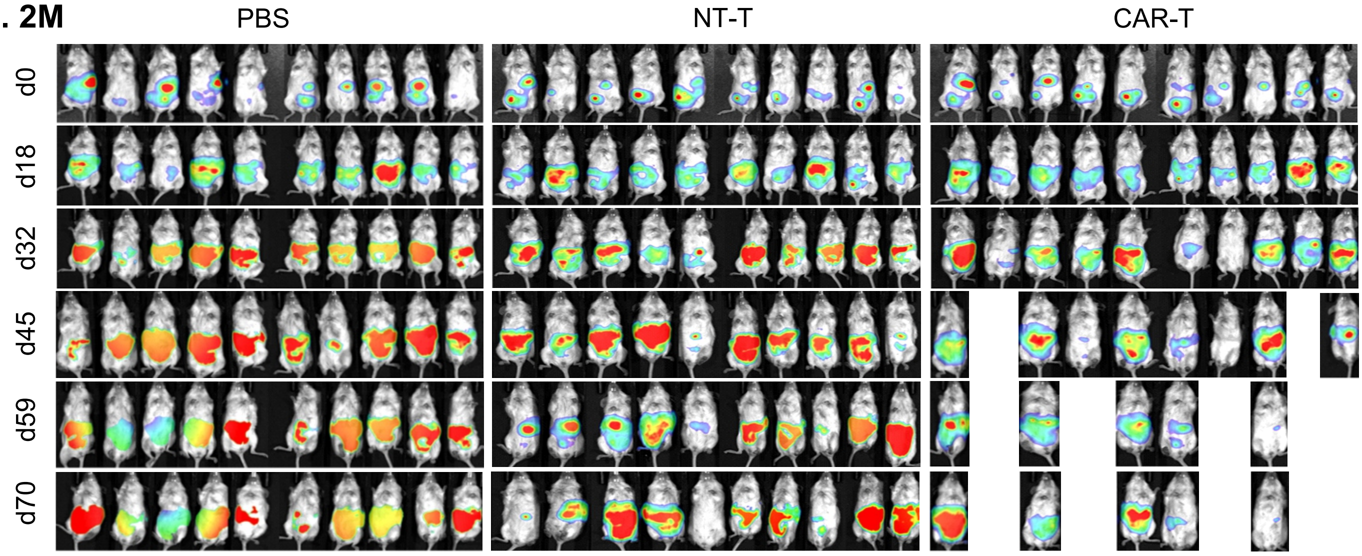
